# Single extracellular vesicle imaging and computational analysis identifies inherent architectural heterogeneity

**DOI:** 10.1101/2023.12.11.571132

**Authors:** Kshipra S. Kapoor, Seoyun Kong, Hikaru Sugimoto, Wenhua Guo, Vivek Boominathan, Yi-Lin Chen, Sibani Lisa Biswal, Tanguy Terlier, Kathleen M. McAndrews, Raghu Kalluri

## Abstract

Evaluating the heterogeneity of extracellular vesicles (EVs) is crucial for unraveling their complex actions and biodistribution. Here, we identify consistent architectural heterogeneity of EVs using cryogenic transmission electron microscopy (cryo-TEM) which has an inherent ability to image biological samples without harsh labeling methods and while preserving their native conformation. Imaging EVs isolated using different methodologies from distinct sources such as cancer cells, normal cells, and body fluids, we identify a structural atlas of their dominantly consistent shapes. We identify EV architectural attributes by utilizing a segmentation neural network model. In total, 7,576 individual EVs were imaged and quantified by our computational pipeline. Across all 7,576 independent EVs, the average eccentricity was 0.5366, and the average equivalent diameter was 132.43 nm. The architectural heterogeneity was consistent across all sources of EVs, independent of purification techniques, and compromised of single spherical (S. Spherical), rod-like or tubular, and double shapes. This study will serve as a reference foundation for high-resolution EV images and offer insights into their potential biological impact.

## Introduction

EVs are universally released by both prokaryotic and eukaryotic organisms^1, 2^. EVs are encompassed by a membrane that mirrors the characteristics of the originating cell’s plasma membrane-a lipid bilayer that exposes extracellular domains containing proteins and carbohydrates^3, 4^. During biogenesis, EVs undergo the orchestrated incorporation of an extensive spectrum of molecular species, encompassing lipids, proteins and nucleic acids, either actively or passively^5, 6^. As conveyors of cellular information to the surrounding milieu, EVs harbor substantial potential in innovative diagnosis^2, 7, 8^. Furthermore, they are also under investigation for their utility in delivering therapeutic cargoes^9–11^ to cells or tissues, capitalizing on their inherent tissue-targeting capabilities^12^. The exhaustive characterization of the physicochemical and biomolecular constituents of EVs is imperative for their effective utilization in drug delivery applications^10^. Physicochemical properties such as the shape and chirality of synthetic nanoparticles is indicated to play a pivotal role in determining their interactions with cells and tissues^13–15^. Morphological variations in synthetic nanoparticles can impact their distribution patterns within tissue and cells, potentially influenced by the rate of uptake and accumulation within organelles like endosomes^13, 16–22^; however, it is unclear whether such morphological variability also exists in EVs. The prevailing bulk ensemble methods utilized for characterizing EVs, such as western blotting or proteomics technologies, have facilitated the identification of proteins associated with EVs^23^. However, these ensemble averaging techniques tend to mask rather than unveil the inherent heterogeneity of EVs. Recent advancements in single-vesicle analysis^24^ have begun to surmount such limitations and pave the way to decipher EV heterogeneity.

EVs share many of the physical and chemical properties and biogenesis pathways of viruses^25^. In this regard, a virus infecting an individual cell has the capability to generate progeny exhibiting diverse morphologies^26^. This diversity is advantageous for a given virus, enhancing its probability of evading immune responses and antiviral interventions^26–29^. Drawing inspiration from the diversity in architectural shapes of pathogenic viruses^13, 30^, we undertook a systematic investigation to analyze and comprehend the spectrum of morphological variations inherent to EVs. Deciphering the structural characteristics of EVs is pivotal in understanding the mechanisms through which they might execute their multifaceted biological roles.^24, 31, 32^

In order to unravel morphological variability at single-vesicle level, we established a pipeline utilizing cryogenic transmission electron microscopy (cryo–TEM) imaging which enables visualization of the structure of biological molecules, such as proteins and viruses, at high resolution^33^. One of the major advantages of cryo-TEM is that it allows for the visualization of biological molecules in their native state, without chemical fixation or staining^33, 34^. This is particularly useful for studying biological particles such as EVs or viruses, as the chemical treatments used in traditional electron microscopy can alter their structure and thereby hinder their true architecture. Over recent years, the development of synthetic and biological nanoparticles have expanded into a broad range of clinical applications. To facilitate the robust clinical translation of such promising synthetic and bio nano-enabled technologies it is becoming imperative to report the shape and size at a single-particle level for such nanoparticles^22, 35^. Therefore, we established a label-free method: to image; perform high-quality biological segmentation using a custom pre-trained neural network model; quantify and classify single EVs purified from a diverse set of samples. We report here a large dataset of cryo-TEM EV images. Our study reveals substantial morphological diversity among individual EVs – irrespective of the source of origin and isolation method – this diversity is governed as an innate EV physicochemical property and not due to isolation, source, or storage artifacts.

## Results

### A pipeline for visualization of native structures of EVs by cryo-TEM

To determine the structures of EVs, we performed cryo-TEM on EVs that were isolated from tumorigenic cells, non-tumorigenic cells, and serum obtained from healthy donors. We used three different EV isolation techniques: size exclusion chromatography (SEC); differential ultracentrifugation (dUC); density gradient (DG) (**Figure 1A**). Initially, the isolated EVs are vitrified on the EM grids and then imaged using cryo-TEM (**Figure 1B, C**). This technique allows for excellent preservation of the specimen in a near-native environment: this allowed us to accurately measure morphological parameters of the EVs. We generated 701 micrograph datasets with sufficient resolution to identify the EVs. To handle the large imaging datasets generated, we established a segmentation neural network model for image data processing and informatics workflow using Cell Pose 2.0, an open-source state-of-the-art biological image analysis tool^36^ (**Figure 1D-E**).

**Figure. 1.**
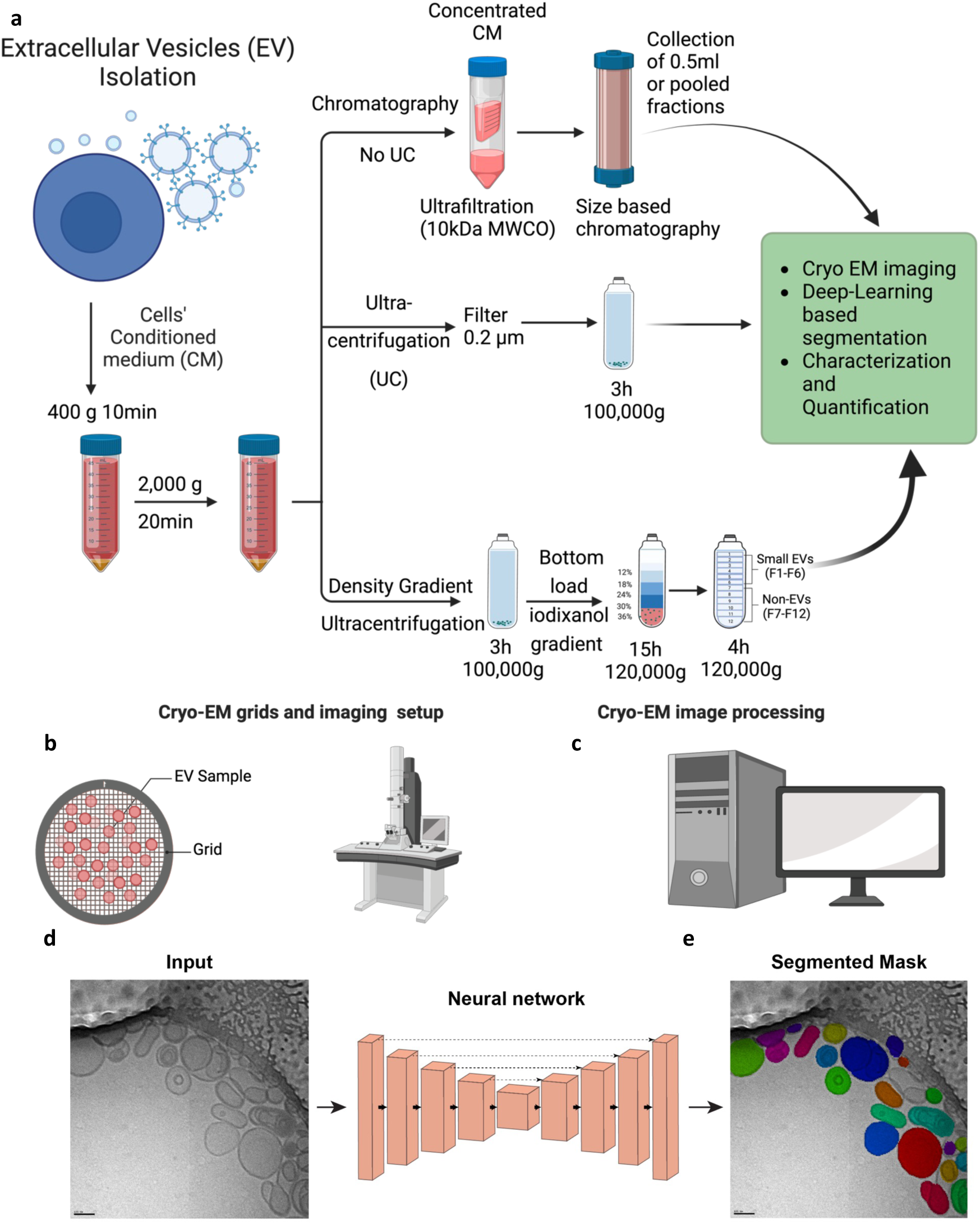
A pipeline for visualization of native structures of EVs by cryo-TEM. **(A-E)** Schematic overview of the workflow. **(A)** Extracellular vesicles samples are isolated via three different isolation methods. **(B-C)** The EV samples are then applied to the grid and undergo cryo-TEM imaging and processing **(D-E)** A neural network model is trained to perform biological segmentation of extracellular vesicles from cryo-TEM micrographs and obtain a segmented mask. Scale bar, 100nm.

### Systematic and quantitative analysis identifies architectural diversity within EVs

We first imaged EVs derived from tumorigenic (Panc1 and T3M4) and non-tumorigenic (HPNE and HEK293T) cells via differential ultracentrifugation (dUC). A total of 1,503 particles were analyzed for Panc1, T3M4, HPNE and HEK293T EVs isolated via dUC. Most of the visualized particles could be classified into 3 major categories: Single Spherical (S. Spherical); Tubular and Double. All the analyzed particles were classified as EVs due to the clear presence of the lipid bilayer/membrane. The EVs derived from Panc1 cells showed structural diversity (**Figure 2A-E**), consistent with previous predictions^37^. We next investigated EVs from additional cell lines (HPNE, T3M4 and HEK293T) and observed similar structural variability across all EVs, irrespective of their cellular source (**Figure 2F-J, Supplementary Figure 1A-E, Supplementary Figure 2A-E**).

**Figure. 2.**
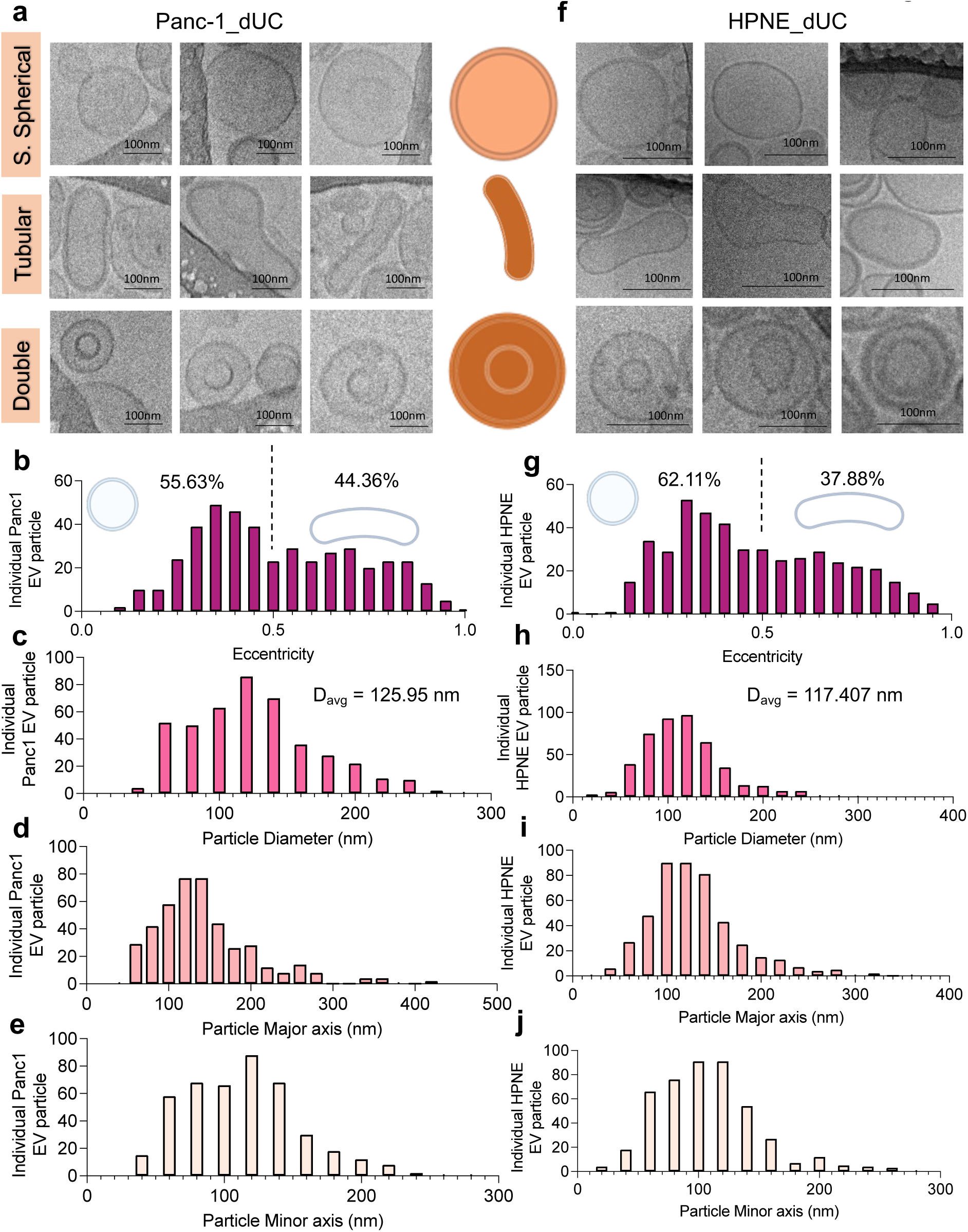
Systematic and quantitative analysis identifies architectural diversity within EVs isolated from Panc1 and HPNE cells via dUC. **(A, F)** Three example Panc1 and HPNE dUC isolated extracellular vesicles for each category are shown. Individual EV particle **(B, G)** eccentricity, **(C, H)** particle diameter, the average particle diameter is indicated, **(D, I)** particle major axis and **(E, J)** particle minor axis parameters are quantified. N = 435 independent dUC-Panc1 EVs and N = 454 independent dUC-HPNE EVs were imaged and quantified for 3 biological replicates. Scale bars denoted on respective image.

Eccentricity was used as a quantitative metric for defining the shape of individual EVs. When eccentricity is close to 0, the particle is considered a perfect sphere. The range of eccentricity is 0 ≤ e < 1. When e is 0.5, an EV is considered more elliptical and when e > 0.5, the EV is considered rod-like or tubular. For Panc1 dUC purified EVs, 55.63% of individual EVs had low eccentricity (e < 0.5), revealing that such EVs were spherical to elliptical in nature. The average equivalent diameter of Panc1 and HPNE dUC purified EVs was 125.95 nm and 117.407 nm, respectively (**Figure 2C, H**), within the characteristic size range of exosomes^1^. The next category of the EVs were the tubular, which were distinguished by having a rod-like architecture (**Figure 2A, F**). EVs were quantitatively grouped into tubular category when the eccentricity of individual EV was greater than 0.5. We found that both Panc1 dUC and HPNE dUC EVs contained tubular EVs, albeit at a lower frequency than S. spherical and double EVs (**Figure 2B-G**). The third classification category was the double vesicles (**Figure 2A, F**), which are smaller EVs encapsulated within a larger EV with more spherical morphology. Double EVs exhibited similar spherical morphology like the single spherical EV category. Double EVs contributed to the percentage of EV particles having low eccentricity (0<e<0.5).

### Validation of varying morphologies of EVs using different isolation techniques

Distinct methods for isolating the EVs offer unique strengths and limitations concerning various factors such as the isolation principle, purity and yield^38–40^. Our objective was to explore whether the structural variability of EVs, encompassing the different categories we listed, remains consistent across various isolation methods (**Figure 3A**). Our aim was to investigate whether the morphological diversity identified in the vesicles are indeed intrinsic components of the EVs and not subject to bias or a result of an artifact caused by specific methods of EV isolation. Thus, in addition to using dUC, we isolated EVs from four cell lines (Panc1, T3M4, HPNE and HEK293T) using size exclusion chromatography (SEC) without any involvement of high-speed ultracentrifugation and two cell lines (Panc1 and HPNE) using OptiPrep-based density gradient (DG) (**Figure 3A**). In the DG method, EVs were loaded at the bottom layer of the gradient and subsequently purified based on their inherent buoyant floatation density^41^. For SEC method, the conditioned media (CM) was first concentrated using an ultrafilter. The concentrated conditioned media (CCM) was then loaded onto a qEV size-exclusion column, and 24 fractions were collected. The EVs were found to elute predominantly in fractions 7 to 10, as verified by quantitative protein analysis, nanoparticle tracking analysis and immunoblotting for the EV markers syntenin-1 and CD81 and the exclusion marker histone H3 (**Supplementary Figure 3A-D**).

**Figure. 3.**
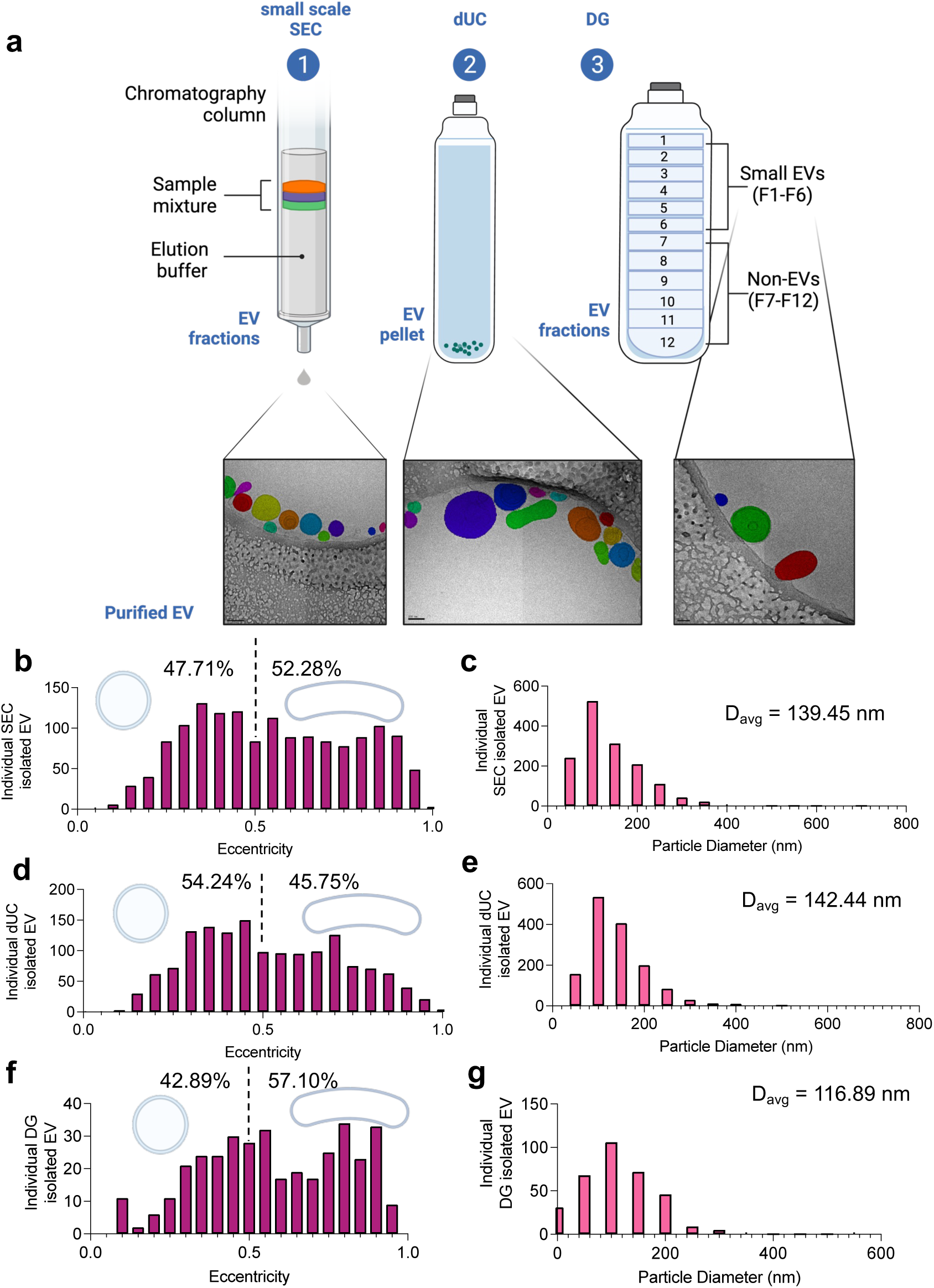
Validation of varying morphologies of EVs using different isolation techniques. **(A)** Isolation of EVs via three isolation techniques size exclusion chromatography (SEC), differential ultracentrifugation (dUC) and density gradient (DG) with representative images in the insert. A total of 3,375 independent cell line derived EVs were imaged via cryo-TEM using three distinct isolation methods and individual EV **(B-G)** eccentricity and diameter of respective isolation technique are quantified. Scale bars denoted on respective image.

Across all the cell source EVs imaged via cryo-TEM using three distinct isolation methods a total of 3,375 independent single-vesicles were quantified (**Figure 3B-G**). We observed that EVs isolated by different cell sources via distinct isolation techniques resulted in similar eccentricity distribution enriching for EVs that were both spherical and tubular in morphology (**Figure 3B, D, F**). The average equivalent diameter of all cell source derived EVs via SEC, dUC and DG are 139.45 nm, 142.44 nm and 116.89 nm respectively (**Figure 3C, E, G**). The major and minor axes for EVs isolated from three isolation techniques are compared in (**Supplementary Figure 4A-F**). Next the independent quantification of eccentricity, the equivalent particle diameter and the major and minor axes of all n = 1,506 individual EVs quantified across all four cell lines and via SEC isolation method are showcased in (**Figure 4A-J**; **Supplementary Figure 5A-E** and **Supplementary Figure 6A-E**). Quantification of eccentricity, the equivalent particle diameter and the major and minor axes of all EVs quantified across the two cell lines and via DG isolation method (n = 366 individual EVs) are showcased in (**Supplementary Figure 7A-E**, **Supplementary Figure 8A-E**).

**Figure. 4.**
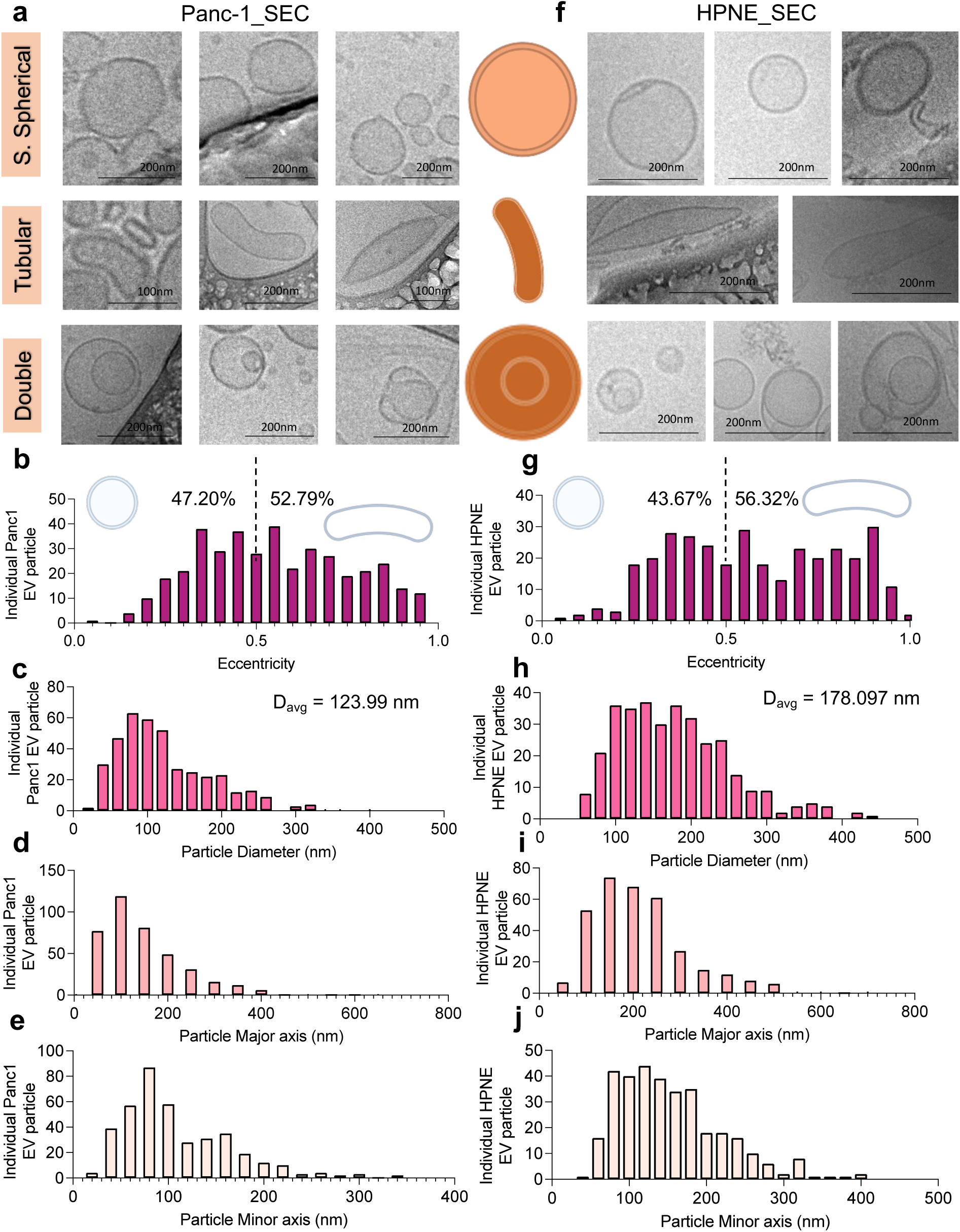
Systematic and quantitative analysis identifies architectural diversity within EVs isolated from Panc1 and HPNE cells via SEC. **(A, F)** Three example Panc1 and HPNE SEC isolated extracellular vesicles for each category are shown. Individual EV particle **(B, G)** eccentricity, **(C, H)** particle diameter, the average particle diameter is indicated, **(D, I)** particle major axis and **(E, J)** particle minor axis parameters are quantified. N = 394 independent SEC-Panc1 EVs and N = 332 independent SEC-HPNE EVs were imaged and quantified across 3 biological replicates. Scale bars denoted on respective image.

**Figure. 5.**
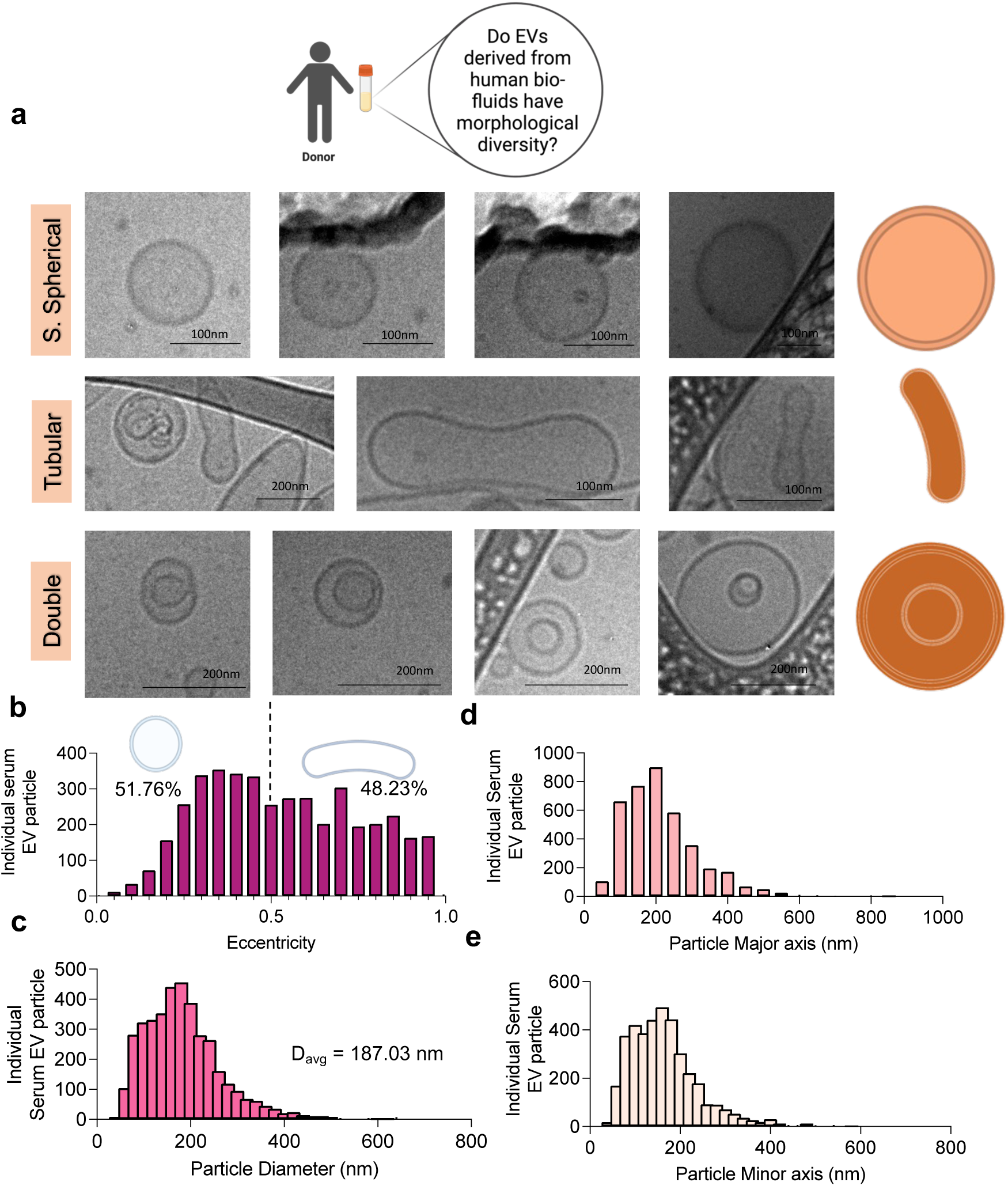
Variable morphologies are conserved in EVs derived from human biofluids. **(A)** Examples of serum isolated extracellular vesicles for each category are shown. Individual EV particle **(B)** eccentricity, **(C)** particle diameter, the average particle diameter is indicated, **(D)** particle major axis and **(E)** particle minor axis parameters are quantified across n = 4171 independent serum EVs from n=3 healthy individual donors. Scale bars denoted on respective image.

**Figure. 6.**
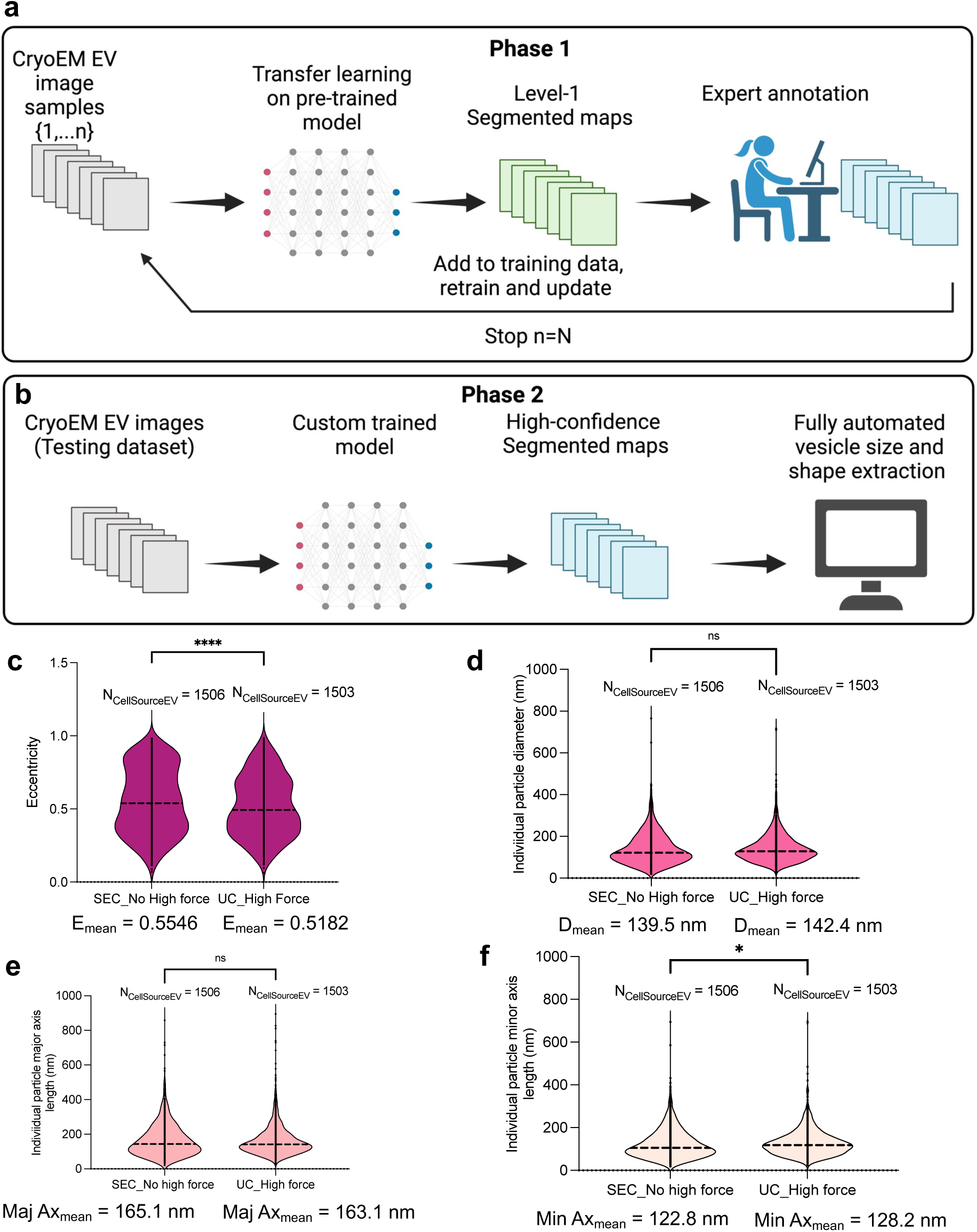
Pre-trained neural network model for quantifying structural characteristics of EVs. **(A, B)** Schematic of phase 1 and phase 2 standardized image data processing and informatics workflow for performing high-confidence segmentation of EVs from cryo-TEM micrographs. During phase-1 segmentation masks are created from scratch on the training dataset. These phase-1 segmentation masks are used as starting point for correction in the expert annotation step. As more images are corrected the model corrected maps are added to the training dataset and the model is re-trained. During phase-2, the custom trained model is run, and high confidence EV-segmented maps are generated. Computationally comparing **(C)** eccentricity, **(D)** EV diameter, **(E)** EV major and **(F)** minor axis, for EVs that were subjected to high force (ultracentrifugation steps; n = 435 dUC-Panc1 EVs, n = 454 dUC-HPNE EVs, n = 485 dUC-T3M4 EVs, n = 129 dUC-HEK293T EVs; N_Total_ = 1503 independent EVs, across three biological replicates) versus EVs that were not subjected to high force (No ultracentrifugation steps; n = 394 SEC-Panc1 EVs, n = 332 SEC-HPNE EVs, n = 321 SEC-T3M4 EVs, n = 459 SEC-HEK293T EVs; N_Total_ = 1506 independent EVs, across three biological replicates). Short-dashed line represents the median of the datapoints in the Violin plots. Statistical analysis was determined by using unpaired two-tailed t-test and **** denoted p < 0.0001, * denoted p = 0.022 and ns: not significant.

### Variable morphologies are conserved in EVs from biofluids

To further validate if these structural variabilities are observed in EVs derived from biofluids, we isolated EVs from human serum samples. Cryo-TEM analysis revealed the presence of all three major sub-categories (S. Single, Double and Tubular) in all the samples analyzed (**Figure 5A-E**). In EVs derived from biofluids, 51.76% of independent EV particles had low eccentricity suggesting the enrichment of a more spherical population. These data suggest that the morphological diversity is conserved in biofluids.

Next, keeping in mind the potential membrane damage that could be caused due to sample storage at-80°C, we tested if there is variability in morphologies for EVs that are freshly isolated versus EVs that were stored at-80°C and subjected to a freeze/thaw cycle. In comparison to the freshly prepared EVs, freezing (for 30 days) and subsequent thawing of EVs did not influence their morphological variability (Supplementary Figure 9A-D). There was insignificant difference observed between the equivalent particle diameters of fresh versus EVs stored EVs, thereby making storage artifacts less likely to be a contributing factor for observed structural variabilities. In literature, morphological diversity is observed among EVs from plasma, breast milk and cerebrospinal fluid (CSF)^42–44^. Taken together, these results suggest that EV structural diversity is identified in EVs purified from human biofluids.

### Pre-trained neural network model for quantifying structural characteristics of EVs

To quantify the large imaging datasets of EVs we established a standardized image data processing and informatics workflow (**Figure 6A-B**). The workflow consisted of two stages: (1) a robust segmentation of extracellular vesicle particles, and (2) structural characteristics extraction of the segmented particles. For the segmentation stage we used a state-of-the-art deep neural network model adapted from Cellpose 1.0^45^ and Cellpose 2.0^36^. Once segmented, the physical characteristics are extracted using in-built image processing algorithms that measure region properties of labelled segments.

Cellpose^36^ is a cellular segmentation algorithm powered by a deep learning neural network, whose architecture is a modified U-net style architecture with residual blocks^46, 47^. Cellpose provides different pre-trained models that are trained on segmentation datasets of cytoplasm and nuclei. However, using these pre-trained models on our EV data produced sub-par results. To improve the performance, in phase 1 (**Figure 6A**), we used an expert-in-the-loop approach to transfer-learn a custom model by training on a small subset of our data. We started with the pre-trained model called ‘cyto2’ and iteratively feed-in images from our cryo-EM data. At each output, an expert corrected the level-1 segmentation by manual annotation. These updated maps, post expert annotation were then added back to the training data and the model was retained and updated (**Figure 6A**).

Once desirable segmentation maps were achieved, the training was stopped and phase 2 of testing was done using our custom-trained deep-neural network model (**Figure 6B**). Our training dataset utilized a small number of training images (n=32), and the rest of the n=669 images were used as the testing dataset to generate high-confidence EV segmentation maps. Once high-confidence EV segmentation maps were achieved, the maps were passed through a fully automated algorithm for EV size and shape extraction (**Figure 6B**). We found that without our fine-tuning and the expert annotation approach the segmented vesicle image generated by the pre-trained neural network model was performing undesired segmentation of the sample grid elements while missing substantial number of EVs (Supplementary Figure 10A-F).

Using the model graphical user interface (GUI) we were able to correct the model’s mistakes on a single image by drawing the region of interest (ROI) that were missed or eliminating the ROIs that were incorrectly picked up by the model. Using this corrected EV segmented image as the ground-truth annotation, a new Cellpose model was trained (we name the model EVpose) and applied to the next image from our dataset. We observed that after retraining the model with only 32 images from our dataset of 701 images, the model’s accuracy of segmenting the EVs reached 95% (**Supplementary Figure 10D-F**). We stopped the iterative process after 32 images, once we were satisfied with the segmentation accuracy and then utilized the rest of the image dataset (n=669 images) as the testing dataset.

After achieving the satisfactory individual EV segmented masks, we fed these masks into second stage algorithm that determines physical parameters such as single vesicular: i) diameter, ii) eccentricity, and iii) major and minor axis of individual EVs in the cryo-EM micrography. Here, we define eccentricity (e) as a parameter that directly determines the shape of the EVs. For EVs that belonged to a perfect spherical morphology its eccentricity was close to 0 and EVs that had a more rod-like or tubular architecture the eccentricity was close to 1. Using our model, we were also able to compute and extract the size (equivalent particle diameter) and shape (eccentricity) metrics for EVs that were subjected to no force (no high-speed centrifugation) while isolation versus EVs that were subjected to high force (high speed centrifugation) during EVs isolation (**Figure 6C-F**). We show through this high-throughput quantitative analysis that across all n=7,576 individual vesicles the average eccentricity was found to be close to 0.5366 (**Supplementary Figure 11A-D**), suggesting most of the EVs represented an elliptical architecture irrespective to the mode of EV isolation and the source of EV formulation.

Overall, our model system needed only about 40 minutes to run and finish generating vesicle segmented masks from our testing dataset (n=669 images); database save the quantitative metrics for all the 7,576 individual extracellular vesicles identified via cryo-EM analysis and store the mask overlayed images on data servers. The entire training and testing process required only 1-2 hours of user time for EVpose as opposed to potentially weeks of work needed with current traditional methods (such as ImageJ^48^) for quantification for such a large image dataset.

### Phosphatidylethanolamine (PE) may contribute to the morphological heterogeneity of EVs

A major question that remains unaddressed is how these morphological variabilities occur in the EVs and what could be the reasons for EV membrane curving/remodeling to occur. Here, we provide a speculation: i) We found via an in-silico analysis from the ExoCarta database^49, 50^ that there is presence of BAR (Bin/Amphiphysin/Rvs) domain family proteins enriched in EVs (List of BAR domain family proteins found in EVs is listed in Supplementary Table 1). It has been shown previously that membrane associated BAR proteins and phospholipids, play a role in shaping the cell membrane^51–53^. ii) Phospholipid ions such as phosphatidylethanolamine (PE), DAG, Ceramides, phosphoinositides (PI) and cholesterol are major influencers in shaping the biological membrane curvature and structure^51, 52^. Owing to the role of phospholipid in regulating the shape and curvature of lipid membranes, we determined whether PE was also found in EVs and whether this could be contributing factor for architectural variability and determine EV membrane curvature. We identify via positive high mass resolution spectra that PE specific ions were associated with normal and PDAC cell derived EVs (**Supplementary Figure 12A-D**). We hope future studies will further probe such speculations with functional experiments.

## Discussion

Cryogenic transmission electron microscopy (cryo-TEM) is one of the electron microscopy methodologies employed for characterizing EVs, with this technique the risk of sample damage and the introduction of artifact effects are mitigated but yields images with a lower contrast^37^. Cryo-TEM involves rapid freezing, commonly utilizing liquid ethane^54, 55^, whereby water vitrifies instead of forming orderly crystals, thereby preserving the native architectural and structural properties of EVs^56^. Cryo-TEM facilitated the initial visualization of exosomes in 2008^57^, and by capturing membrane bilayers and EV structures, cryo-TEM has the potential to unravel EV morphology^37, 42, 54, 58, 59^. However, systematic examination of morphological variability of EVs isolated from multiple sources (normal cells, cancer cells, serum) and using distinct isolation methods along with a highly effective, rapid and fully automated workflow to quantify cryo-TEM micrographs are still lacking and crucial for standardized analysis of large datasets.

To fully quantify and appreciate the architectural diversity of individual extracellular vesicles, a large number of single EVs need to be imaged and analyzed. Here, we analyzed the morphological variability across different EV samples using cryo-TEM imaging approach. Using this approach, we identified 7,576 individual EVs and quantified specific parameters such as particle eccentricity, diameter, major and minor axis of individual vesicles. Our data suggests that EVs belonging to the single spherical category is the most abundant morphological category associated with EVs derived from different sources.

We observed that the traditional ‘cup-shaped’ EV morphology imaged via negative stain EM is likely an artifact caused by the sample preparation. Our data suggests that EVs from different cell types have distinct morphological variabilities when imaged via cryo-TEM. The wide-ranging structural variabilities observed in EVs hints at the presence of distinct subpopulations, each potentially characterized by unique functionalities and biochemical attributes. For all the individual EVs (n = 7,576) imaged and quantified, we found that the median eccentricity (e) was 0.5366, suggesting that in the natural native state EVs are not perfectly spherical. Apart from eccentricity, we quantified the individual particle diameter across all analyzed EVs and the average diameter was 132.43 nm, suggesting our different methods of isolation were able to enrich for the small EV population. We further quantified the major and minor axes of individual EVs, which were 153.82 nm and 118.001 nm, respectively.

Another contribution of our study was to provide researchers with a quick, fully automated platform for quantifying EVs from Cryo-EM micrographs. Analyzing a large dataset of images manually can be very tedious, requiring manual analysis and is extremely time consuming^60^. Therefore, in this study we describe a very rapid, highly efficient, fully automated imaging analysis model for quantitative evaluation of EVs from Cryo-EM micrographs. Our vesicle segmentation approach is based on a pre-trained neural network model with a ‘human-in-the-loop’ approach adapted from the Cellpose 2.0 architecture^36^, and we found our approach succeeded across a wide range of scenarios. Our analysis showed that we were able to train our segmentation model with a very limited dataset (n = 32 images as training set versus n = 669 images as testing set). The pipeline designed in this study can be effectively employed for the segmentation and quantification of a broad spectrum of individual vesicular particles, extending even to larger EVs.

It will be informative to develop methods that would enable the enrichment of vesicles based on their phenotypical diversity (compositional and morphological diversity) to better understand the versatile implications of composition-specific and shape-specific EVs in the development of therapeutics and diagnostics and propel further exploration and address a wide array of biological questions.

## Materials and Methods

### Cell culture

The cell lines employed in our study (HEK 293T, HPNE, Panc1) were obtained from American Type Culture Collection (ATCC); (T3M4) was obtained from the cell bank at RIKEN BioResource. HEK 293T were cultured in Dulbecco’s Modified Eagle Medium (DMEM, Corning) supplemented with 10% fetal bovine serum (FBS) and 1% penicillin-streptomycin. T3M4, HPNE and Panc1 were cultured in Roswell Park Memorial Institute Medium (RPMI 1640, Corning) supplemented with 10% FBS and 1% penicillin-streptomycin. Cells were sub-cultured at a 1:5 or 1:10 ratio every 2-3 days, using Trypsin-EDTA (GIBCO 15400054) to remove the adherent cells from the plate. All the cell lines were maintained in humidified cell culture incubators at 37°C and 5% CO_2_. Panc1, T3M4, HPNE and HEK293T cell lines were validated by the Cytogenetics and Cell Authentication Core at MD Anderson. All cell lines were routinely tested for mycoplasma contamination.

### Extracellular Vesicles production

Cultured cells at a confluency of about 80% were thoroughly washed with PBS and subsequently placed in serum-free medium for 48h. The conditioned medium (CM) of the serum-starved cells was harvested and subjected to centrifugation: 400 x g for 10 minutes to pellet cells. Supernatant was centrifuged at 2000 x g for 20 minutes to remove debris and the pellet discarded. Subsequently, the CM was filtered using a 0.2 μm filter (Fisher). After the initial processing, extracellular vesicles were isolated by the methods described below.

### Cell line-derived EV isolation

Three methods were used for EV isolation: size exclusion chromatography (SEC), differential ultracentrifugation (dUC), Optiprep-based density gradient (DG).

### Size exclusion chromatography (SEC)

After filtration the CM was concentrated using Amicon Ultra-15 10kDa filter. 500 μl of concentrated CM (CCM) was overlaid on 70 nm qEV 500 size exclusion columns (Izon, SP1) for separation. A dedicated column was used for each cell line. The column was flushed in between samples using filtered PBS and filtered 20% ethanol. For each EV sample, twenty-four fractions of 500 μl were collected using the Izon automated fraction collector (AFC v1). The fractions F7 to F25 were analyzed by western blot to probe for EV-enriched and non-EV enriched fractions. The EV-rich fractions were F7-F10 and EV-poor fractions were F15-30. Pooled fractions (7-10) and (11-30) were then concentrated using 10 kDa cutoff filters (Amicon Ultra-15, Millipore). Each fraction was subjected to protein concentration measurement, Nanoparticle Tracking Analysis (NTA) and western blot analysis following manufacturer’s instructions. The EV-enriched fractions (F7-F10) were further concentrated using sterilized Amicon Ultra-15 10kDa filter analyzed by high-resolution cryogenic transmission electron microscopy.

### Differential ultracentrifugation (dUC)

For EVs isolation by dUC method, harvested conditioned media was subjected to ultracentrifugation at 100,000 x g for 3 hours at 4°C using a SW32 Ti Beckman Coulter rotor. After ultracentrifugation, the supernatant was discarded, and the EV pellet was resuspended in PBS. For performing cryogenic transmission electron microscopic imaging EVs were centrifuged and washed in PBS at 100,000 x g for 3 hours at 4°C in a SW41 Ti rotor (Beckman Coulter).

### Density gradient (DG)

EVs that were isolated by UC (1 ml) were subjected to a density gradient purification with OptiPrep Medium (Sigma). The OptiPrep gradients were prepared using ice-cold PBS with following percentages and volumes per gradient: 12% (2 ml), 18% (2.5 ml), 24% (2.5 ml), 30% (2.5 ml) and 36% (2.5 ml). EVs were mixed with the OptiPrep medium to a final concentration of 36% for bottom loading in ultracentrifuge tubes. Very gently, the next gradient fractions were gently layered on top in descending order: 30%, 24%, 18% and finally 12%. Gradients were subjected to ultracentrifugation at 120,000 x g for 15 hours. The following day, 12 fractions of 1 ml each were collected (F1 to F12) and each fraction was washed in ice-cold PBS (12-fold dilution) and subjected to ultracentrifugation at 120,000 g for 4 hours. Each fraction (F1 to F12) was resuspended in PBS and the EV rich fractions (F1-F6) were utilized for cryogenic imaging.

### Human serum-derived EV isolation

Serum samples from healthy participants were considered deidentified discarded material and exempt from requiring approval from the Institutional Review Board (IRB) of University of Texas at MD Anderson Center, and informed consent was obtained from all participants. Serum samples were collected from each participant, and relevant participant information can be found in Supplementary Table 2. Each serum sample was centrifuged at 400g for 10 minutes followed by 2000g for 20 minutes at 4°C. The resulting supernatant was then filtered through a 0.22 um pore size syringe filter followed dUC isolation. Post EV isolation the serum EVs were subjected to high resolution cryogenic transmission electron microscopy characterization.

### Nanoparticle tracking analysis

The concentration and the size distribution of the EVs was measured based on their Brownian motion using a Nano Sight LM10 (Malvern), Blue 488 laser and a highly sensitive sCMOS camera. During measurements, temperature was set and kept constant at 25°C. Syringe pump speed was set to 20. For each acquisition, a 90 second delay followed by three captures of 30 second each was employed. The average values of three captures were used to determine the nanoparticle concentration and the mode of the size distribution.

### Western blot

For western blotting analysis EV samples were heated in Laemmli sample buffer 4X (Invitrogen Catalog No. J60015-AD) at 95°C for 10 minutes. EV samples were then loaded on 4-12% precast polyacrylamide mini gels (Invitrogen) for electrophoretic separation of proteins. Protein transfer was performed on methanol-activated polyvinylidene fluoride (PVDF) membranes by Trans-Blot Turbo Transfer system (1704150; Bio-Rad). Post transfer the membranes were blocked in 5% Bovine Serum Albumin (BSA) in Tris-buffered saline (TBS) with 0.1% Tween-20 detergent (TBST) at room temperature for 1 hour. After completion of blocking the membrane was incubated in primary antibody overnight on a shaker (Primary antibodies: Syntenin-1 (Abcam, ab133267 1:2000); CD81 (Santa Cruz, sc-166029 1:1000); Histone H3 (Abcam, ab201456 1:1000)). The next day, secondary antibodies were incubated for 1 hour at room temperature (Secondary antibodies: Anti-Rabbit HRP-conjugated Cell Signaling Technology (CST-#7074S 1:5000); Anti-Mouse HRP-conjugated (R&D, HAF007, 1:5000)). Post primary and secondary antibody incubations the membranes were washed with TBS containing 0.1% Tween-20 on a shaker, three times at 10 minute intervals. The visualization of the membrane was performed with West-Q Pico ECL solution (Gen-depot) following the manufacturer’s instructions. Amersham Hyper film (GE Healthcare) was used for capturing the chemiluminescent signals.

### Cryogenic transmission electron microscopy

EVs were resuspended in 50 μl of 1x PBS with a minimum concentration of 5.5×10^10^ EV/50 μl. Quanti-foil mesh grids and Lacey Carbon only (400 Mesh Cu) grids were glow discharged for 2 minutes just before the vitrification step. During vitrification, approximately 3-5 μl of the EVs samples were applied on to the grids, blot time was optimized for 2-5 seconds, blotting force was optimized between (1∼4) to obtain good-quality EV film and followed by snap freezing into liquid ethane using the FEI Vitrobot Mark IV system. The frozen grids were placed in a cryo specimen holder and then transferred to liquid nitrogen for the examination. The samples were imaged using a JEOL 2010 cryo-TEM (200kV, LaB6 filament), with a Gatan 626 cooling holder operated at-180 °C. Mini Dose System (MDS) were used for image processing. The vitrification and imaging steps were performed at Shared Equipment Authority (SEA), Rice University.

### Computation support specification

The end-to-end model (Cellpose segmentation computation and EV parameter quantification) were performed on Windows 11 OS with AMD Ryzen 5950X 16-Core processor (3.4 GHz). The installed RAM was 64GB (Intel, USA). We used GPU (NVIDIA RTX 4090) for the all the human-in-the-loop experiments. Cellpose 2.0 computation time measurements were also performed on this machine.

### Model and training

Our model consists of two main blocks: 1) segmentation block and 2) quantification block. The Python scripts are written in Python 3.8. The initial script, which manages the segmentation stage, employs Cellpose version 2.0.4, alongside numpy version 1.21.6 and PyTorch 1.11.0. To quantify the segmentation quality of the masks generated by the segmentation algorithm, we matched the predicted mask with a ground-truth mask that was annotated with human-in-the-loop approach. We trained the U-NET model with a total of 32 images from our dataset. For reducing the computation times, we down sampled our image size from 2048×2048 to 256×256. After the desired quality segmentation masks were achieved, we utilized 669 images as the training dataset. After training each individual image that down sampled was up sampled again to 2048×2048 and both raw files and overlayed image with mask were saved. Once the image was passed through the segmentation block – the segmented mask was then fed to the quantification block were various single-vesicle parameters were quantified and stored in a list form. The total run times for all the 669 images to pass both the blocks of our model were orders of magnitude shorter (<40 minutes) compared to the time it takes to do the manual ROI segmentation and quantification.

### ToF-SIMS analysis on extracellular vesicles

Positive and negative high mass resolution spectra were performed using a ToF-SIMS NCS instrument, which combines a TOF-SIMS5 instrument (ION-TOF GmbH, Münster, Germany) and an in-situ Scanning Probe Microscope (NanoScan, Switzerland). A bunched 30 keV Bi3+ ions (with a measured current of 0.2 pA) was used as primary probe for analyzing a field of view of 150 × 150 µm^2^, with a raster of 128 by 2128 pixels and by respecting the static limit of 1.1012 ions/cm^2^ to not damage the surface. A charge compensation with an electron flood gun has been applied during the analysis, as well as an adjustment of the charge effects has been operated using a surface potential. The cycle time was fixed to 200 µs (corresponding to m/z = 0 – 3644 a.m.u mass range). PE specific characteristic ion at C_2_H_6_N^+^ (Mass (u) 44.06); Carbohydrate specific characteristic ion at C_3_H_5_O^+^ (Mass (u) 57.02); DAG specific characteristic ion at C_33_H_59_O_4_^+^ (Mass (u) 519.29); SM specific characteristic ion at C_5_H_12_N^+^ (Mass (u) 86.08) are well known characteristic ion peaks for specific phospholipids^61^.

## Acknowledgements

This work was supported by a gift from Fifth Generation, Inc. (“Love ‘Tito’s’”) and funds from Sid W. Richardson Foundation to RK. EV work in the Kalluri lab is supported by NIH R35CA263815, NIH P40OD024628 and NIH R01CA231465. We are grateful to the Shared Equipment Authority staff at Rice University for the support. ToF-SIMS analysis was carried out with support provided by the National Science Foundation CBET-1626418. Graphical figures in this work were created using BioRender.

## Author contributions

Raghu Kalluri: Idea and conceptualization, Study design, Funding acquisition, Project oversight, Resources, Supervision, Writing – Original draft

Kshipra S Kapoor: Co-conceptualization and Ideation, Study design, data curation, formal analysis, investigation, methodology, software, validation, visualization, Writing – original draft

Seoyun Kong: Investigation – assisted KSK

Hikaru Sugimoto: Investigation, Methodology

Wenhua Guo: Investigation, Methodology

Vivek Boominathan: Software, Generation of the neural network model

Yi-Lin Chen: Investigation, Methodology

Sibani Lisa Biswal: Investigation and Resources

Tanguy Terlier: Investigation, Methodology

Kathleen M McAndrews: Investigation, Methodology

---

**Supplementary Figure. 1.**
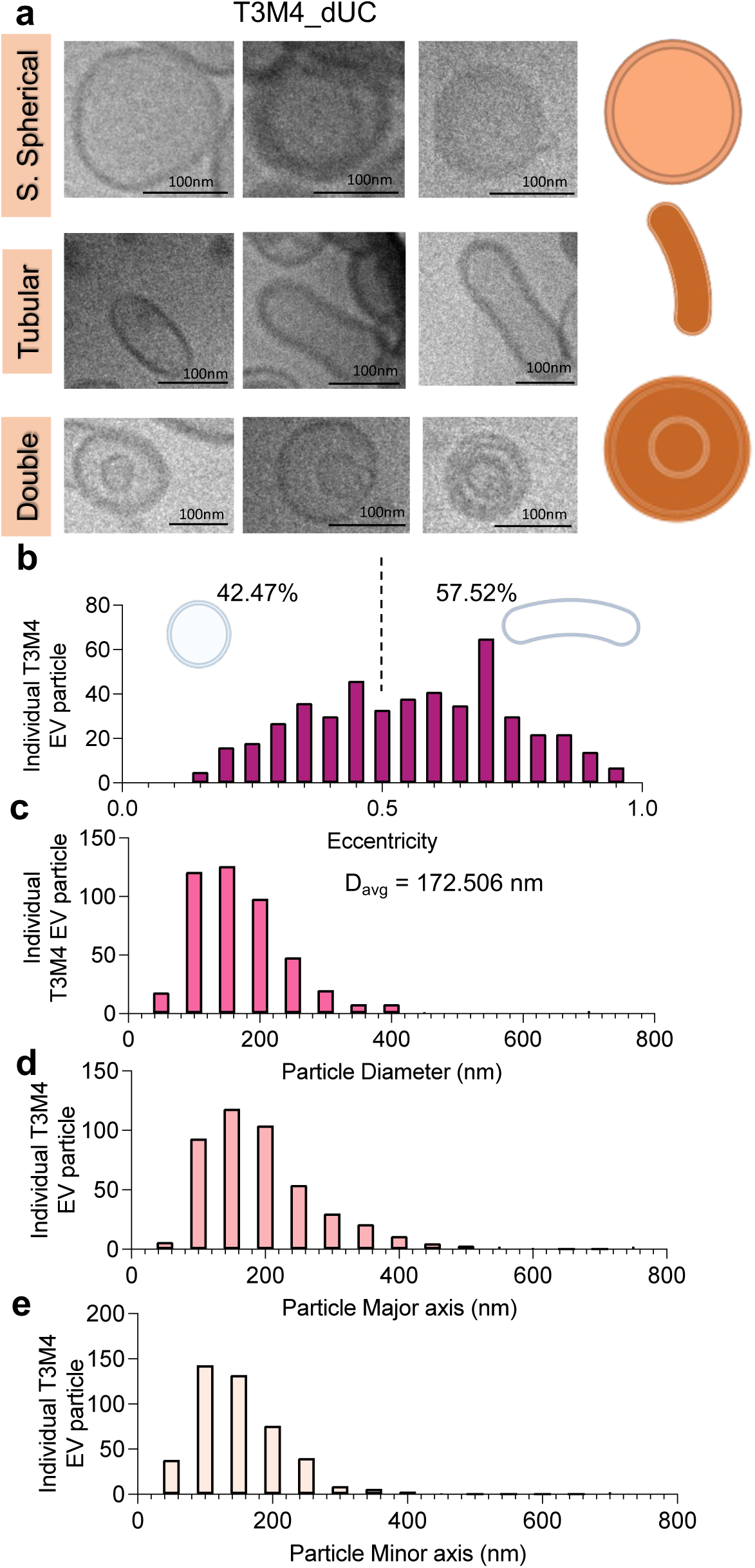
Systematic and quantitative analysis identifies architectural diversity within EVs isolated from T3M4 cells via dUC. **(A)** Three example T3M4 dUC isolated extracellular vesicles for each category are shown. Individual EV particle **(B)** eccentricity, **(C)** particle diameter, the average particle diameter is indicated, **(D)** particle major axis and **(E)** particle minor axis parameters are quantified. N = 485 independent dUC-T3M4 EVs were imaged and quantified, across three biological replicates. Scale bars denoted on respective image.

**Supplementary Figure. 2.**
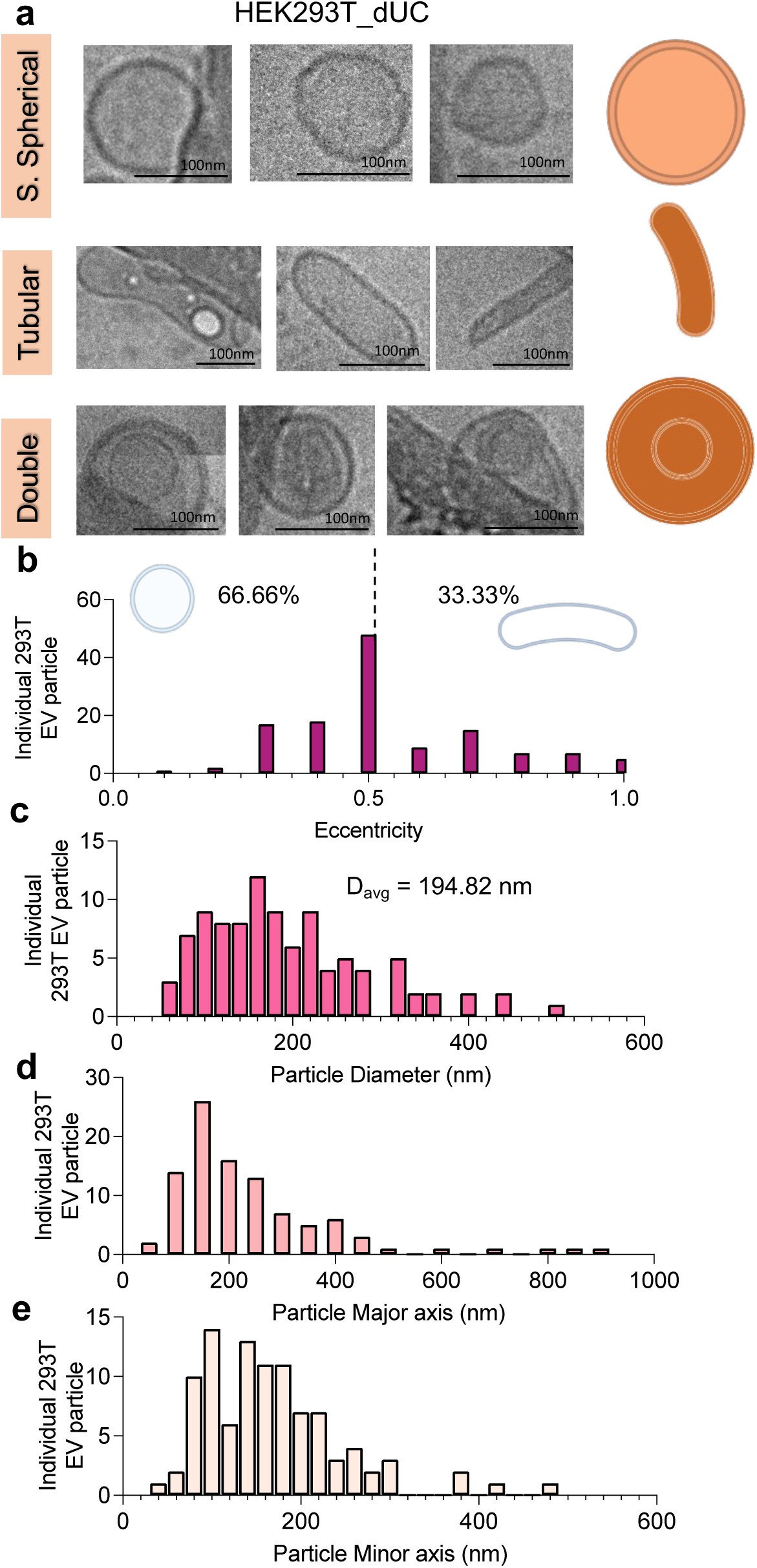
Systematic and quantitative analysis identifies architectural diversity within EVs isolated from HEK293T cells via dUC. **(A)** Three example HEK293T dUC isolated extracellular vesicles for each category are shown. Individual EV particle **(B)** eccentricity, **(C)** particle diameter, the average particle diameter is indicated, **(D)** particle major axis and **(E)** particle minor axis parameters are quantified. N = 129 independent dUC-T3M4 EVs were imaged and quantified, across three biological replicates. Scale bars denoted on respective image.

**Supplementary Figure. 3.**
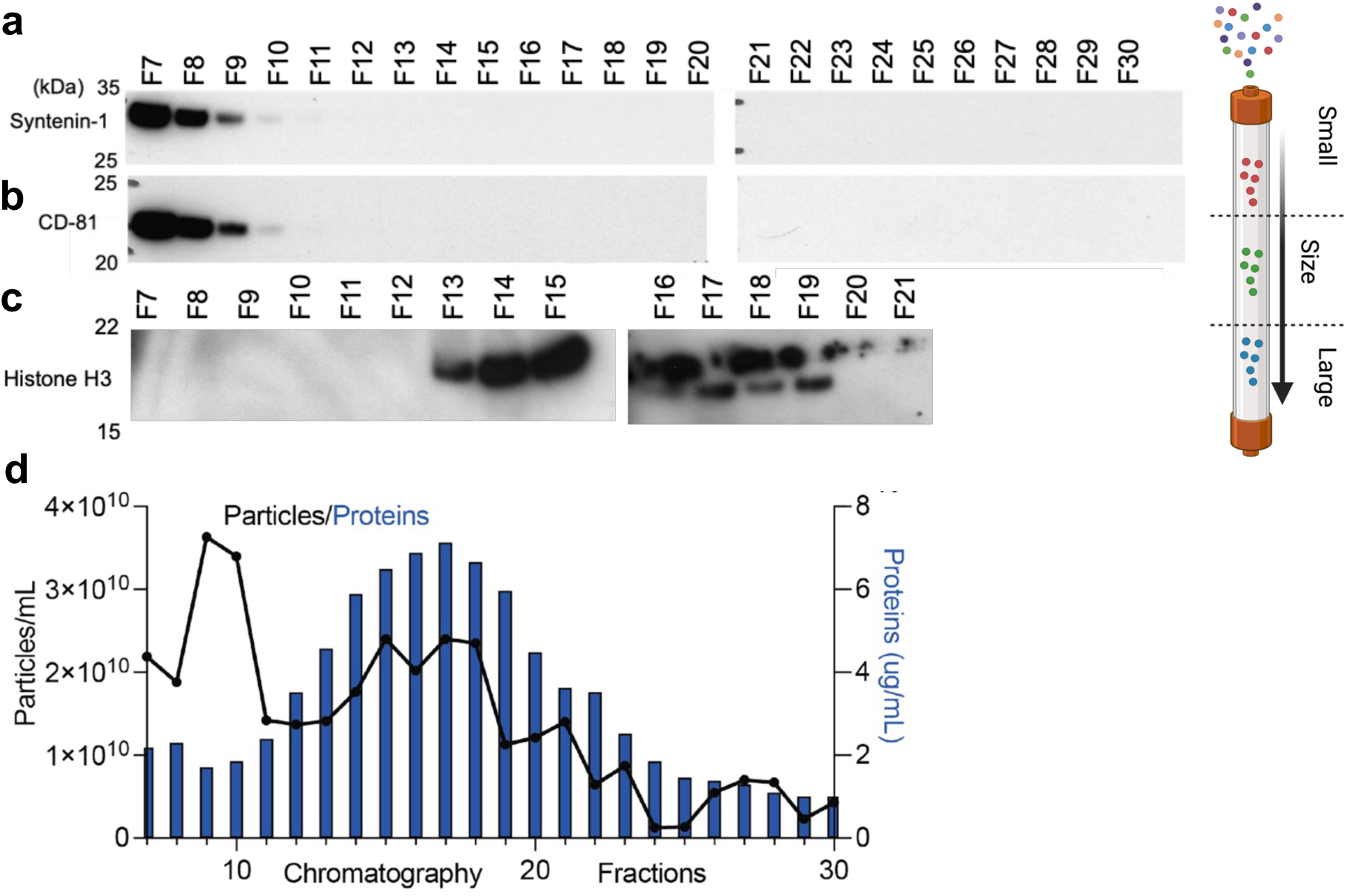
Biomolecular characterization of HEK293T EVs isolated via size exclusion chromatography. **(A-C)** Western blot analysis of EV and non-EV associated proteins of individual SEC fractions. Fractions (F7-F10) are EV-rich fractions and fractions (F12-F22) are protein-enriched fractions. **(D)** Particle count and protein quantification on individual SEC fractions is shown.

**Supplementary Figure. 4.**
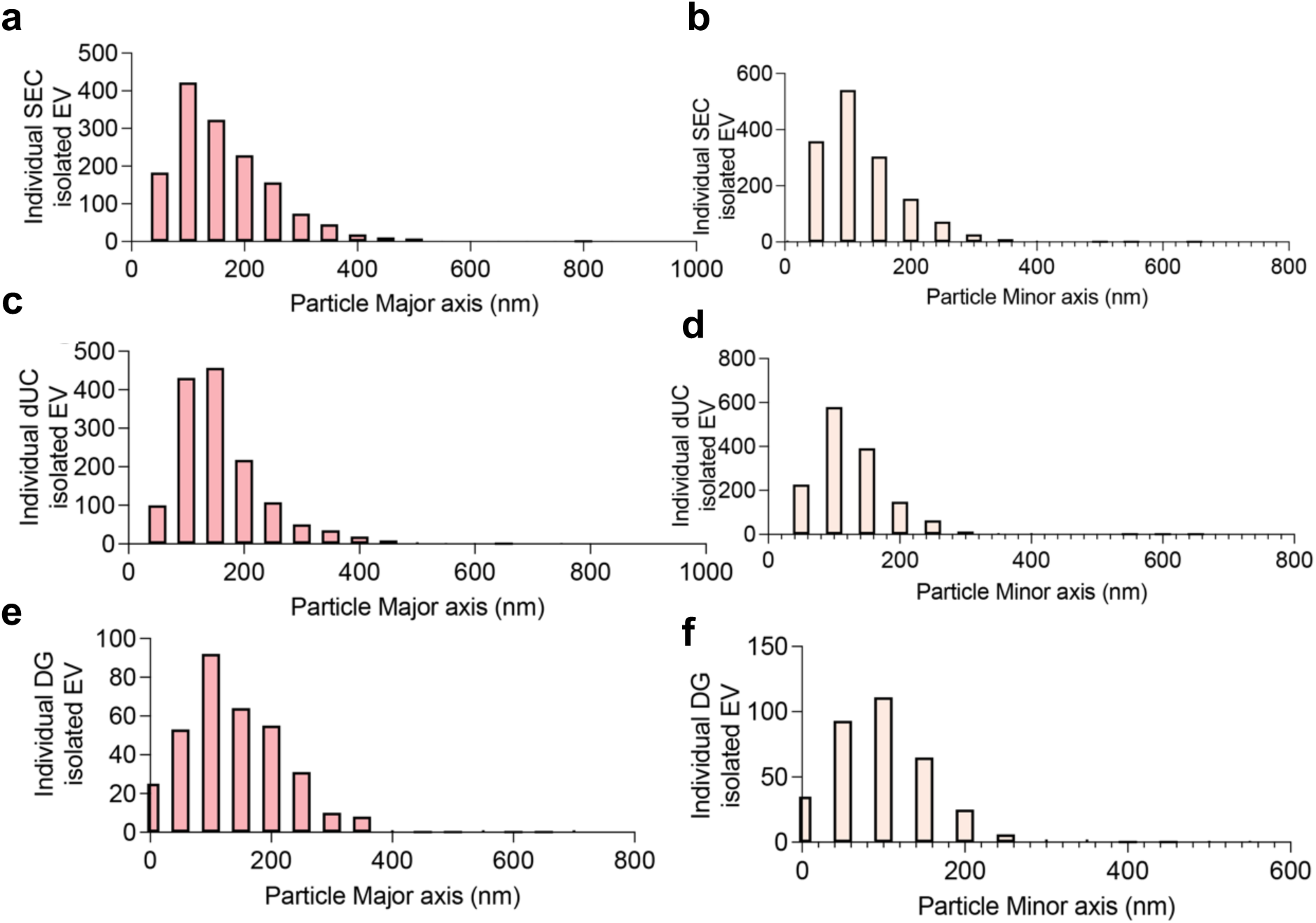
**Quantification of major and minor axis of independent EVs based on different isolation techniques.** (A, B) **SEC** (C, D) **dUC and** (E, F) **DG.**

**Supplementary Figure. 5.**
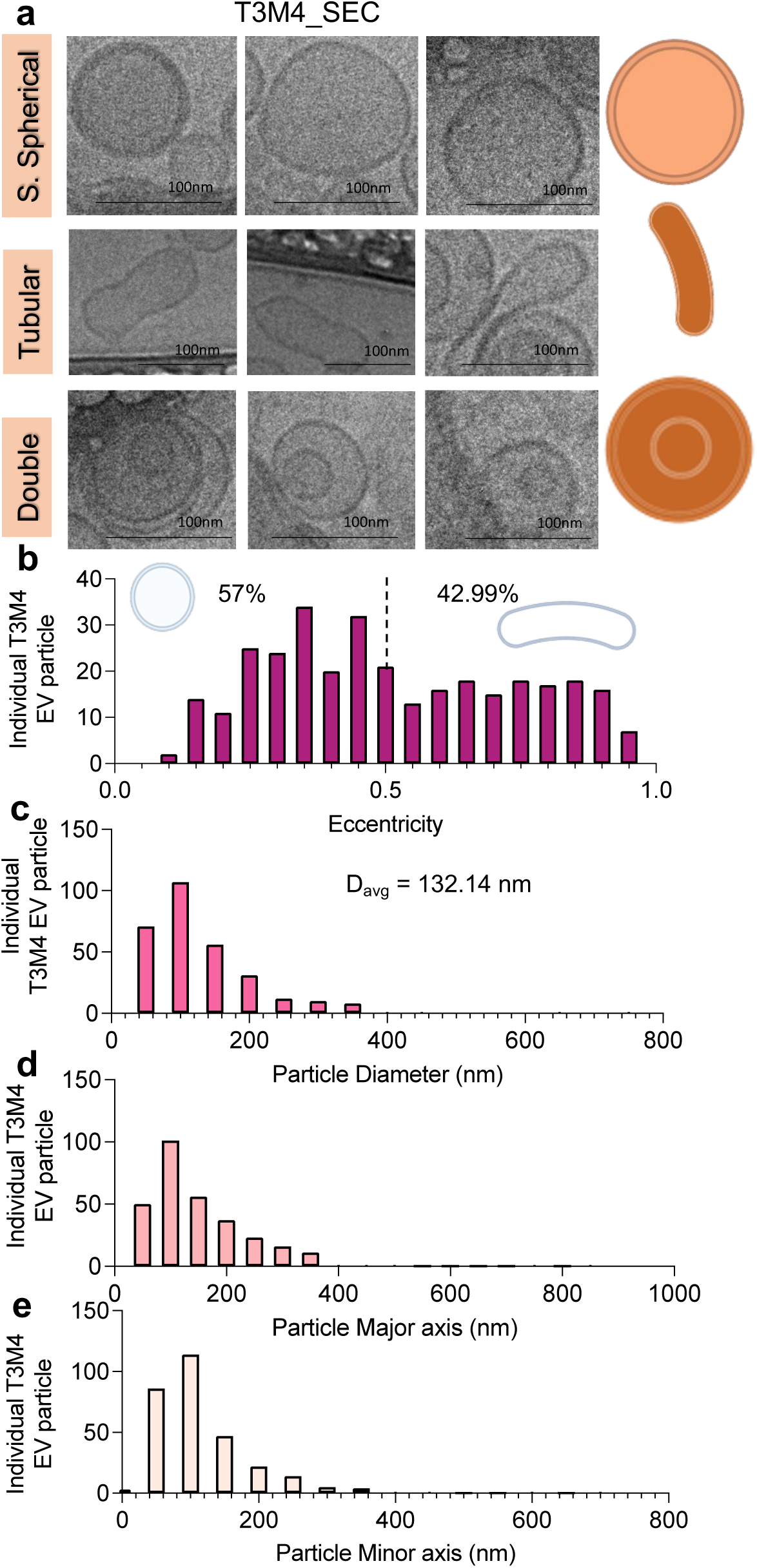
Systematic and quantitative analysis identifies architectural diversity within EVs isolated from T3M4 cells via SEC. **(A)** Three example T3M4 SEC isolated extracellular vesicles for each category are shown. Individual EV particle **(B)** eccentricity, **(C)** particle diameter, the average particle diameter is indicated, **(D)** particle major axis and **(E)** particle minor axis parameters are quantified. N = 321 independent SEC-T3M4 EVs were imaged and quantified, across three biological replicates. Scale bars denoted on respective image.

**Supplementary Figure. 6.**
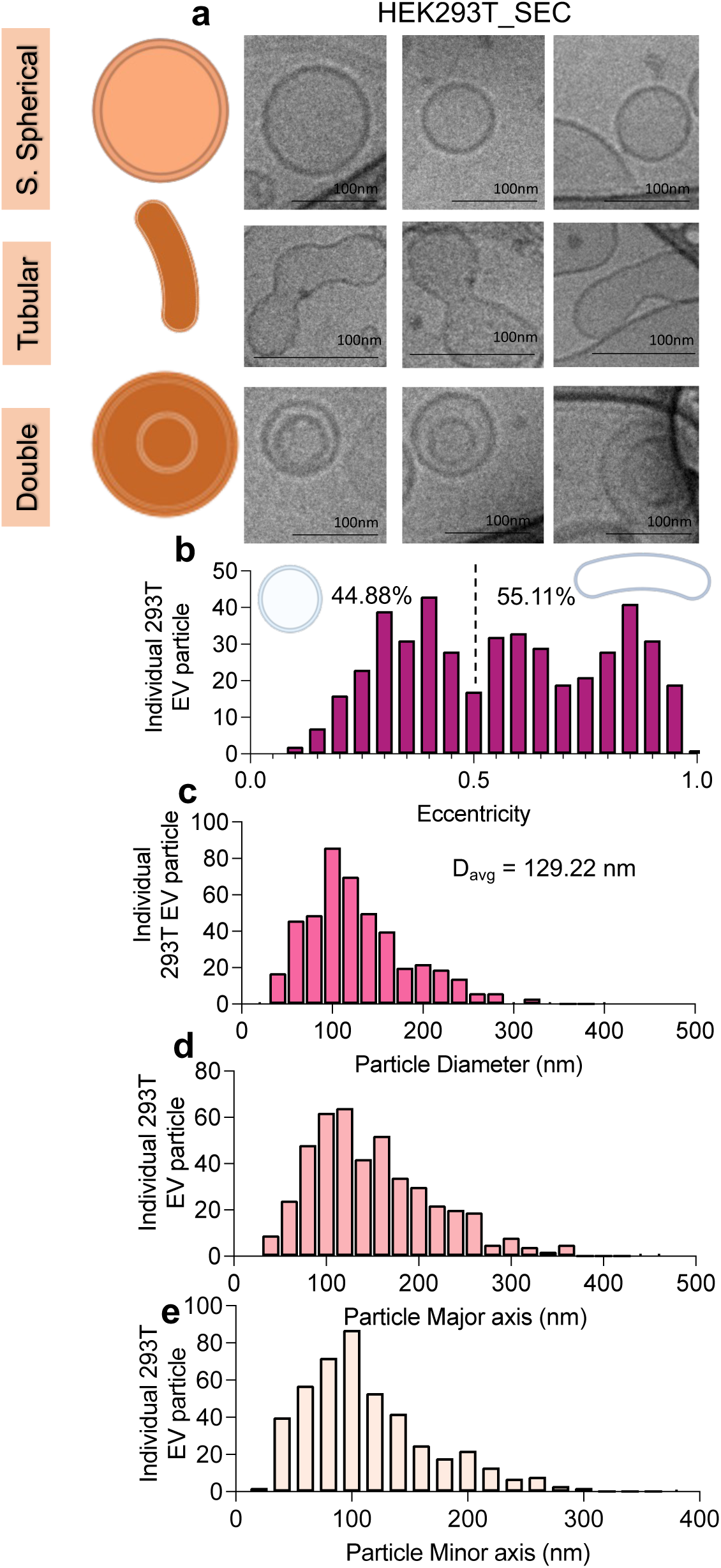
Systematic and quantitative analysis identifies architectural diversity within EVs isolated from HEK293T cells via SEC. **(A)** Three example HEK293T SEC isolated extracellular vesicles for each category are shown. Individual EV particle **(B)** eccentricity, **(C)** particle diameter, the average particle diameter is indicated, **(D)** particle major axis and **(E)** particle minor axis parameters are quantified. N = 459 independent SEC-HEK293T EVs were imaged and quantified, across three biological replicates. Scale bars denoted on respective image.

**Supplementary Figure. 7.**
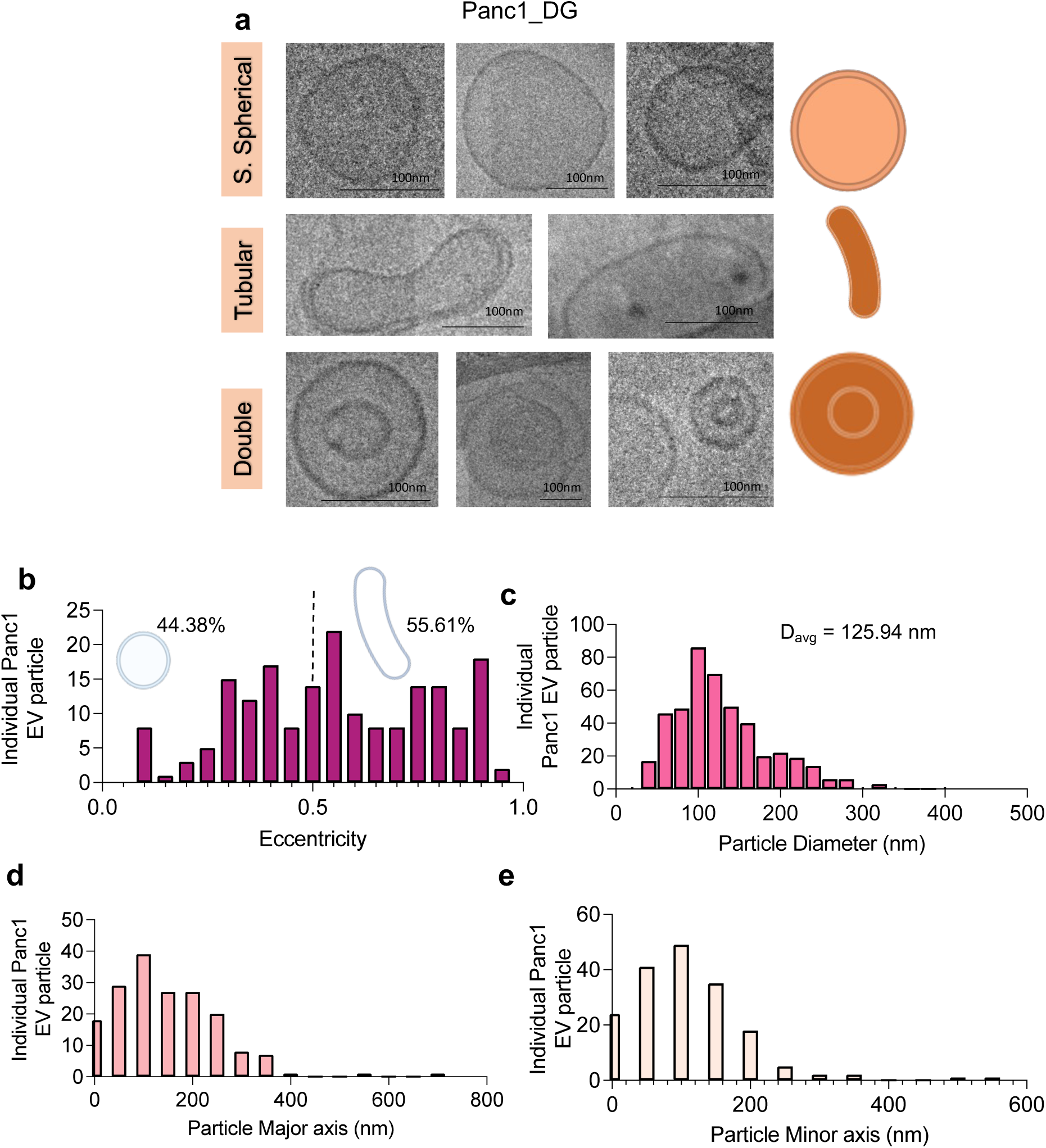
Systematic and quantitative analysis identifies architectural diversity within EVs isolated from Panc1 cells via density gradient. **(A)** Three example Panc1 DG isolated extracellular vesicles for each category are shown. Individual EV particle **(B)** eccentricity, **(C)** particle diameter, the average particle diameter is indicated, **(D)** particle major axis and **(E)** particle minor axis parameters are quantified. N = 187 independent DG-Panc1 EVs were imaged and quantified, across three biological replicates. Scale bars denoted on respective image.

**Supplementary Figure. 8.**
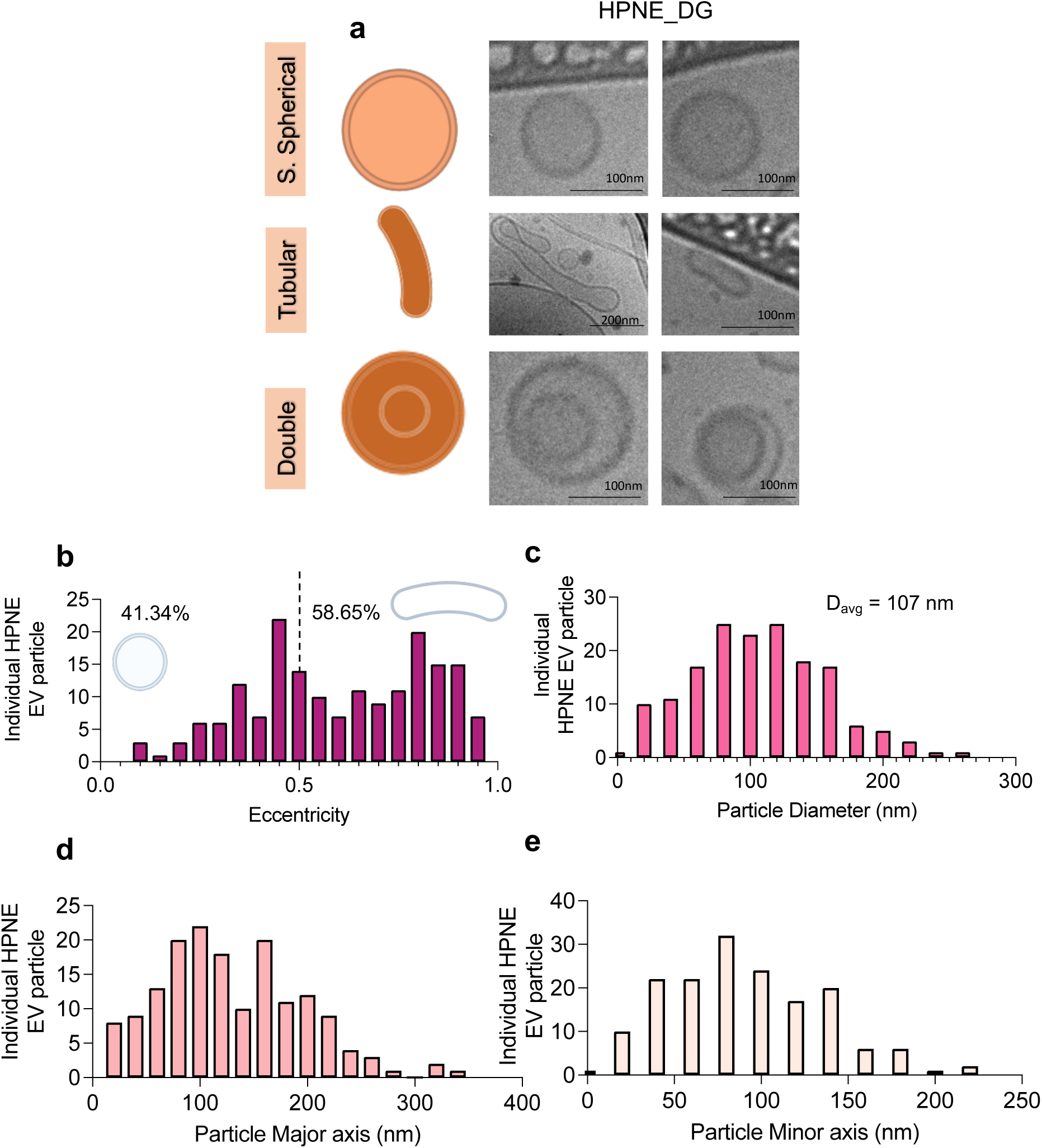
Systematic and quantitative analysis identifies architectural diversity within EVs isolated from HPNE cells via density gradient. **(A)** Two example HPNE DG isolated extracellular vesicles for each category are shown. Individual EV particle **(B)** eccentricity, **(C)** particle diameter, the average particle diameter is indicated, **(D)** particle major axis and **(E)** particle minor axis parameters are quantified. N = 179 independent DG-HPNE EVs were imaged and quantified, across two biological replicates. Scale bars denoted on respective image.

**Supplementary Figure. 9.**
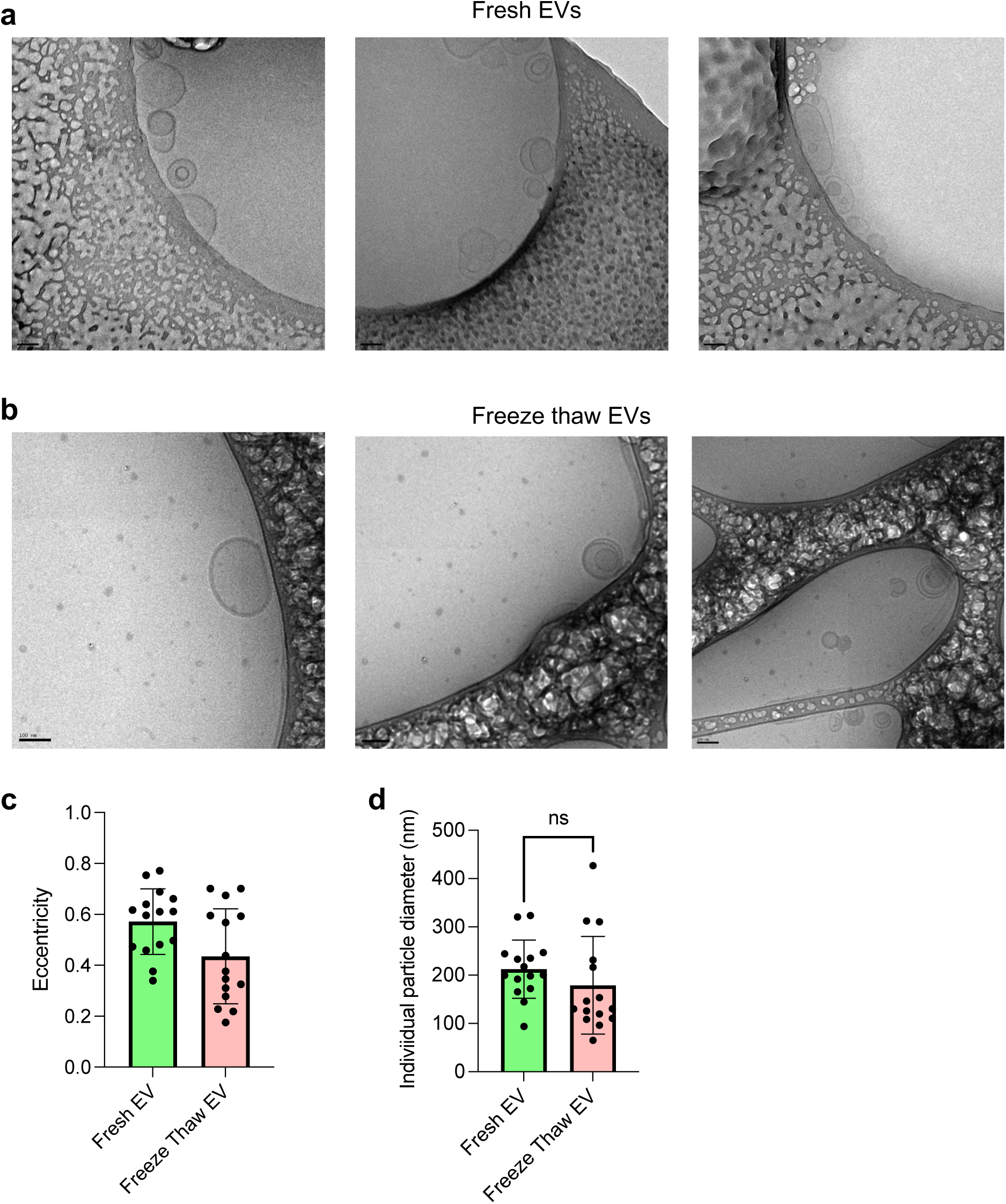
Comparison of EV architecture for fresh EVs versus freeze thaw EVs. **(A, B)** Cryo-TEM images of fresh EV samples – prior to freezing and after freezing (30 days) and thawing. **(C, D)** Comparison of eccentricity and particle diameter for fresh EV samples and EV samples that were subjected to a freeze-thaw cycle (N = 15 independent Panc1 EVs). Data are presented as mean value ± SD. Statistical analysis was determined by using unpaired two-tailed t-test and p = 0.2774. ns: not significant. Scale bars denoted on respective image.

**Supplementary Figure. 10.**
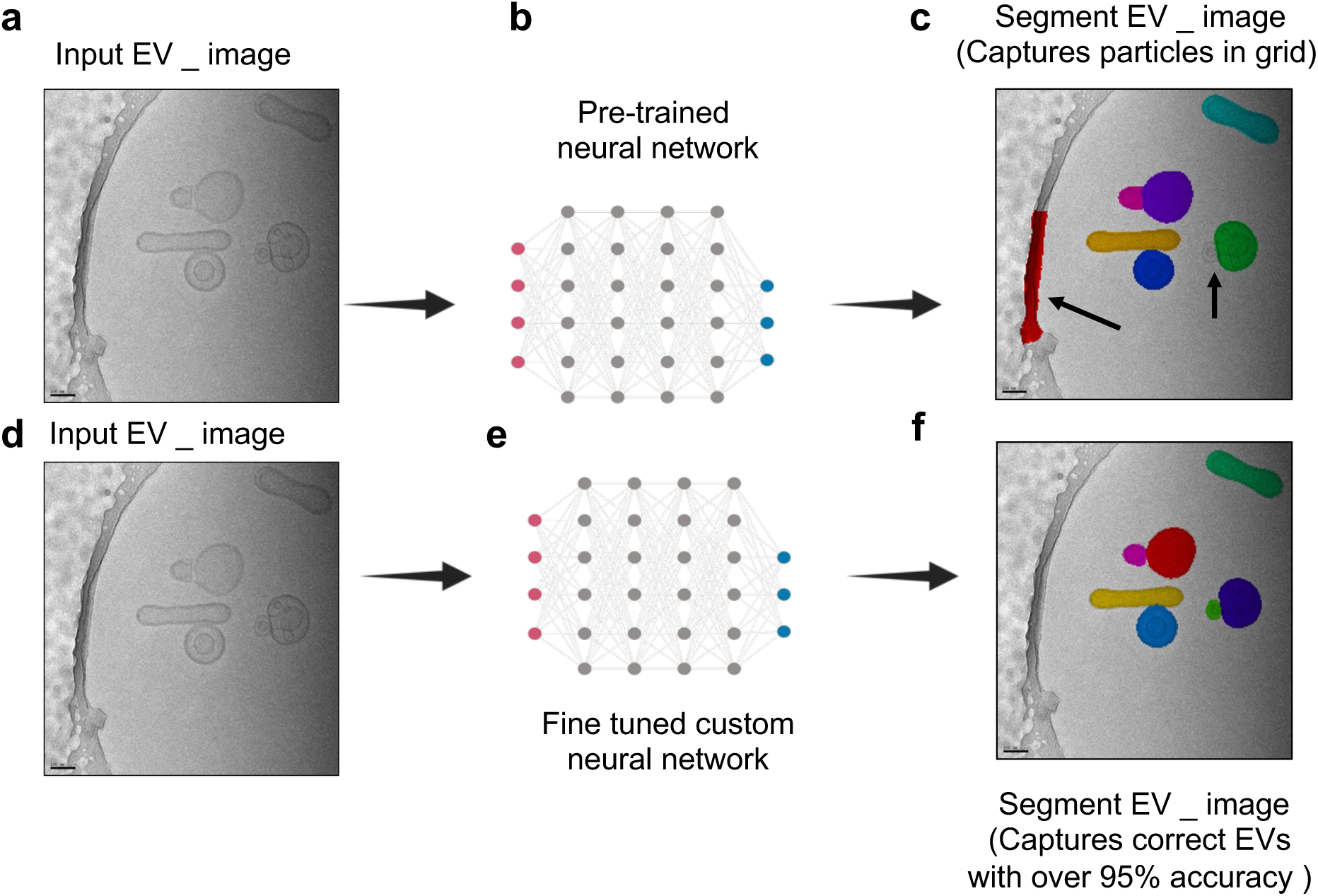
Fine-tuned custom neural network model improved model accuracy for segmenting EVs. **(A-C)** Non-annotated preliminary neural network model mis-interprets and missing EVs in the cryo-TEM micrographs. **(D-F)** Correction by annotation improves the model which thereby decreases the number of errors and achieves high EV-segmentation accuracy. Scale bars denoted on respective image.

**Supplementary Figure. 11.**
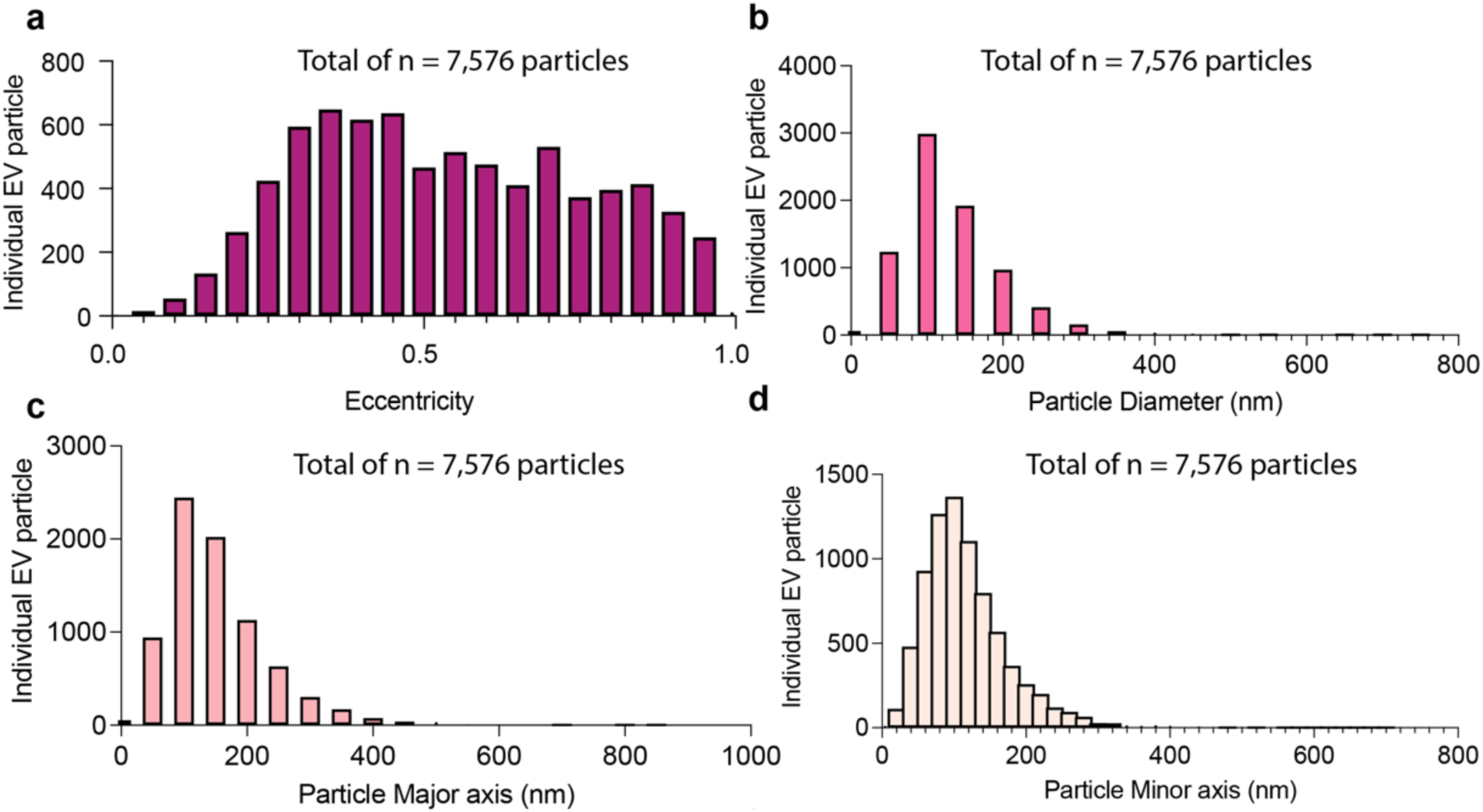
EV analysis of EVs isolated from different sources and isolation techniques. **(A)** Eccentricity, **(B)** Diameter, **(C)** Major axis and **(D)** Minor axis for all 7,576 independent EVs.

**Supplementary Figure. 12.**
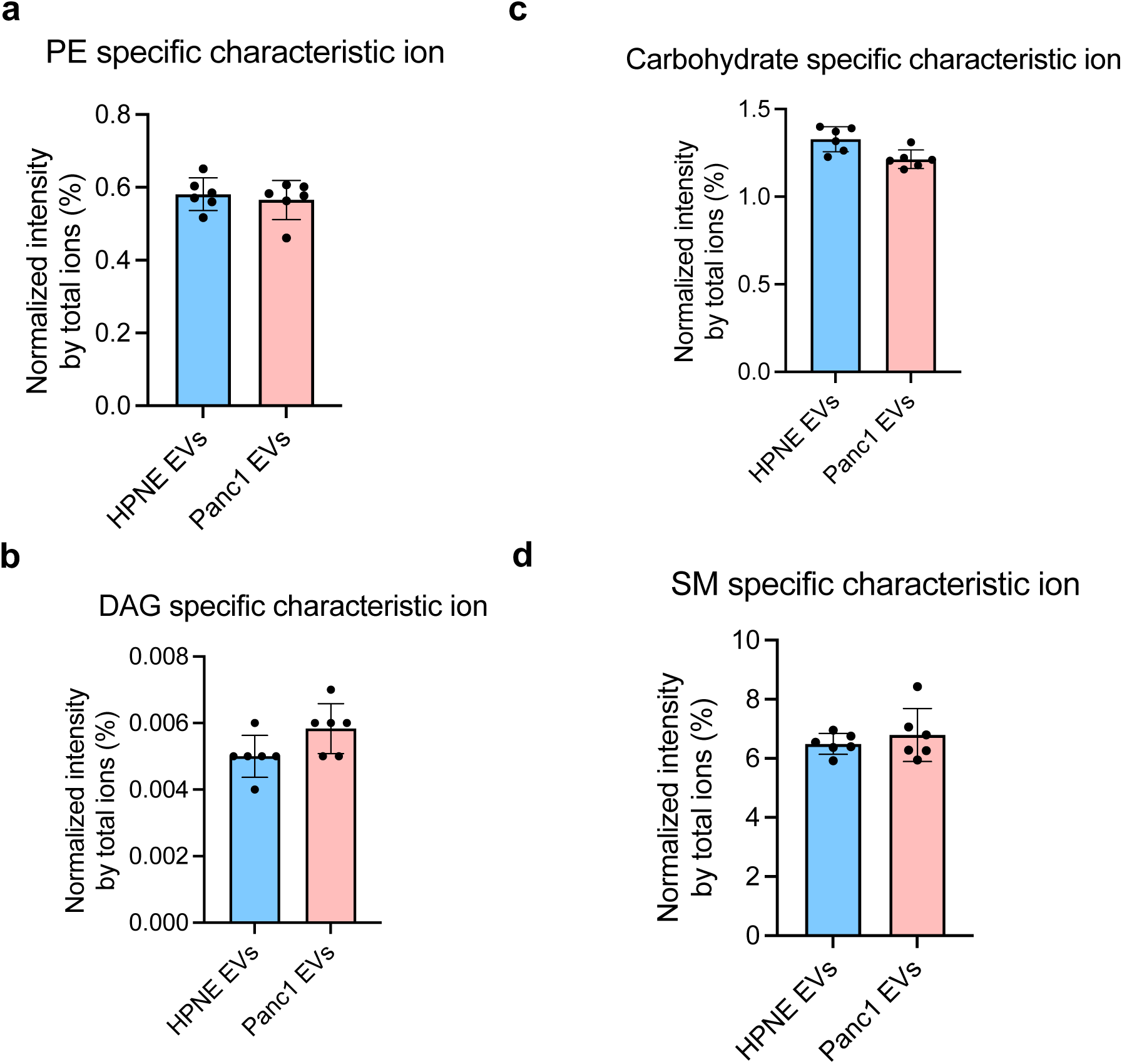
ToF-SIMS analysis to evaluate presence of membrane curving lipids. **(A-D)** ToF-SIMS data for phosphatidyl ethanolamine (PE), carbohydrate, DAG and sphingomyelin (SM) lipid plotted as normalized intensity by total ion intensity for the main peaks (CN^-^, Br^-^). n = 6 technical replicates for HPNE and Panc1 EVs. Data are presented as mean value ± SD.

**Supplementary Table 1.**
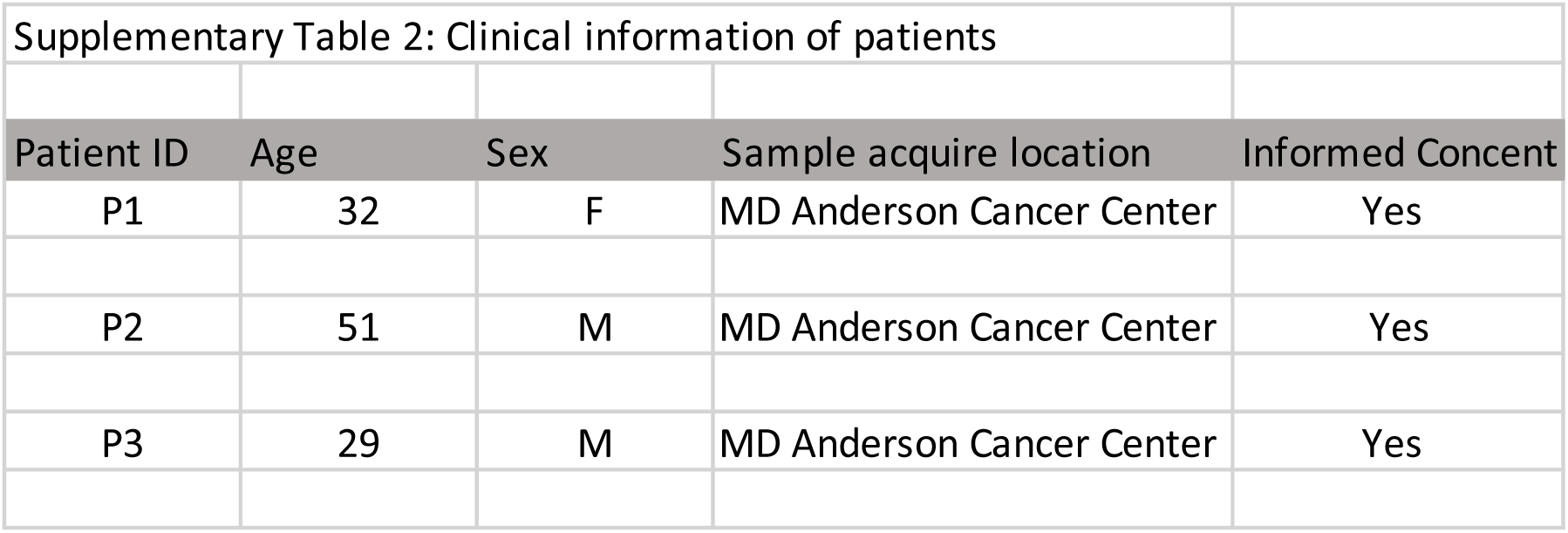

**Supplementary Table 2.**
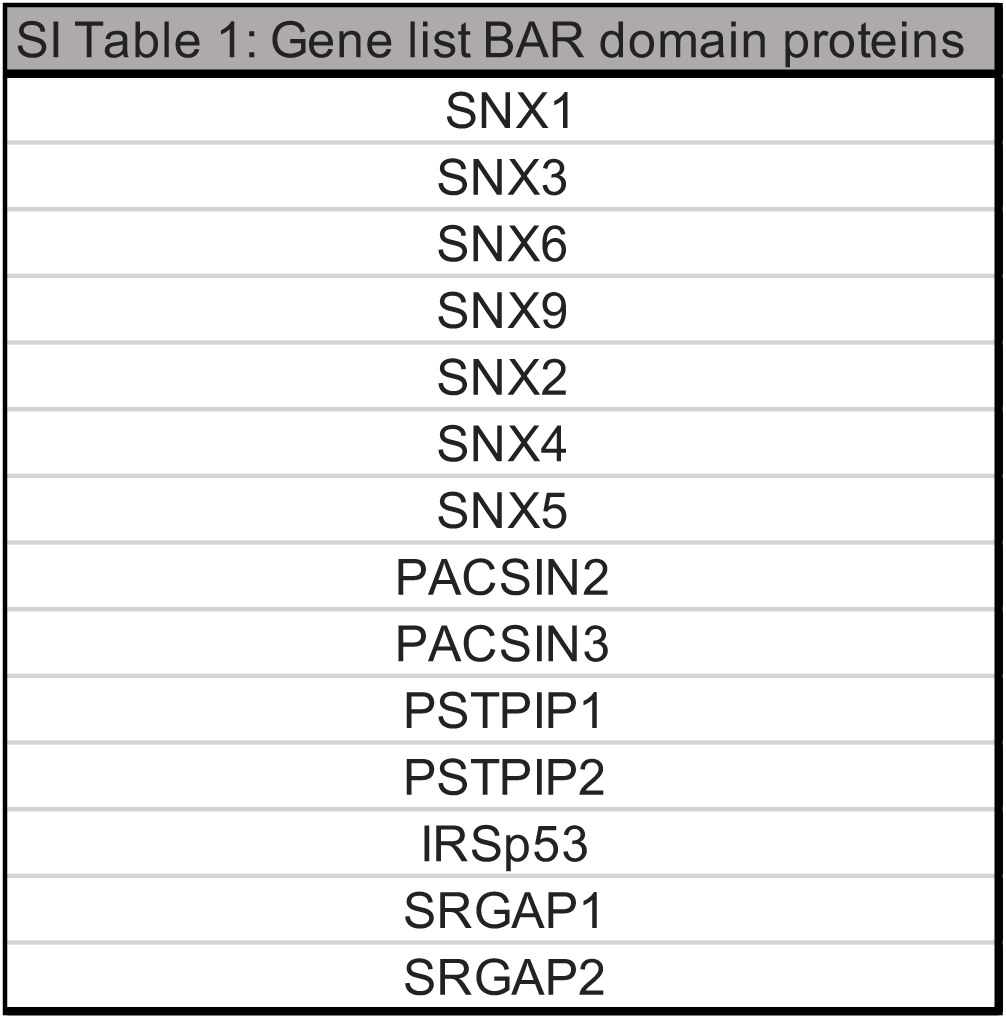

